# Provirus deletion from *Haloferax volcanii* affects motility, stress resistance and CRISPR RNA expression

**DOI:** 10.1101/2024.10.11.617810

**Authors:** Nadia Di Cianni, Simon Bolsinger, Jutta Brendel, Monika Raabe, Sabine König, Laura Mitchell, Thorsten Bischler, Tom Gräfenhan, Clarissa Read, Susanne Erdmann, Thorsten Allers, Paul Walther, Henning Urlaub, Mike Dyall-Smith, Friedhelm Pfeiffer, Anita Marchfelder

## Abstract

*Haloferax volcanii* harbours four putative proviruses: Halfvol1, Halfvol2, Halfvol3 and Halfvol4. In this study we successfully deleted all four provirus genomes, demonstrating, that they are not essential. Transcriptome comparison between this strain (ΔHalfvol1-4) and a wild type strain reveals an increase in archaella and chemotaxis gene expression, resulting in higher swarming motility in ΔHalfvol1-4. Furthermore, ΔHalfvol1-4 cells show an elongated cell shape and a higher resistance to H_2_O_2_ stress compared to the wild type. RNA-seq also revealed down-regulation of CRISPR arrays in the provirus-free strain.

Circularised genomes of Halfvol1, Halfvol2 and Halfvol3 were found in the culture supernatant. This confirms excision of the proviruses from the chromosome, which seems to happen more efficiently at low temperature (30°C). Electron microscopy revealed potential viral particles in the supernatant, and mass spectrometry analysis confirmed the presence of structural viral proteins of Halfvol1 and Halfvol3 in the isolated virus sample. These observations suggest that these proviruses are active and cause a chronic infection in *Hfx. volcanii*.

## Introduction

Viruses are the most abundant entity on the planet and outnumber their microbial hosts by about a factor of 10 (Stern and Sorek, 2010). Given their higher abundance, it is not surprising that cells are commonly found co-infected by multiple viruses, as indicated by many metagenomics studies (Roux *et al*, 2015). Viruses can exhibit different modalities of infection: virulent viruses follow a lytic cycle, resulting in cell lysis, while temperate viruses enter into a latent state, often by integrating their viral DNA into the host genome and being stably inherited. The integrated virus genome is referred to as a provirus and remains latent (or dormant) in a cell population unless induced to excise from the genome and enter a lytic cycle. Spontaneous induction rates are usually very low, but sufficient to allow spread to new hosts. Additionally, some viruses exhibit a chronic cycle, where virus particles are continuously released without lysis of the host cell. Only a limited number of chronic viruses have been documented in prokaryotes, with the majority infecting archaea (Luke et al., 2014; Alarcon-Schumacher et al., 2022).

Some temperate viruses can integrate their genomes randomly into the host chromosome while others integrate in a site-specific manner, frequently into tRNAs (Campbell, 1992). In the case of random integration, a gene may be targeted and disrupted. In the case of site-specific integration, the virus integrates into the host genome by recombining its viral attachment site (*attP*, P for phage) with the host’s attachment site (*attB*, B for bacteria), catalysed by an integrase. As a result of this recombination, the provirus is inserted into the host DNA and flanked by two identical sequences, *attL* (Left) and *attR* (Right). Many archaeal and bacterial temperate viruses integrate into tRNA genes, using part of the tRNA as *attB* (Badel *et al*, 2021; Williams, 2002). We here use the terms *attP* and *attB*, originally established for bacteriophages, also for the archaeal viruses.

Proviruses are surprisingly common in microorganisms (Casjens, 2003; McKerral *et al*, 2023; Munson-McGee *et al*, 2018; Roux *et al*., 2015; Touchon *et al*, 2017), with approximately 60% of microorganism containing at least one functional or defective provirus in their genome (Casjens, 2003). Viruses are generally perceived as enemies of their hosts, engaged in a perpetual arms race with microorganisms. However, proviruses may carry genes that are advantageous for the host itself. The merging of virus and host genomes, even if temporary, creates an ecological window for the evolution of mutually advantageous traits (Feiner *et al*, 2015). Lysogeny thus may promote the development of symbiotic interactions: the long-term association of lysogenic viruses with their host may result in mutually beneficial interactions that enhance the reproductive success of both the virus and its host organism (Feiner *et al*., 2015).

The provirus encoded superinfection exclusion mechanism is an example of such an advantageous trait, and is present in a wide variety of bacterial viruses, such as HK97, T4, D3, P1 and λ (Bondy-Denomy & Davidson, 2014). Superinfection exclusion is a mechanism where a virus which already has infected a host cell prevents subsequent infections by the same or other viruses (Bondy-Denomy & Davidson, 2014; Folimonova, 2012; Hunter & Fusco, 2022). More specifically, superinfection exclusion occurs when a provirus blocks the adsorption (prophage P1), DNA injection (prophage Tuc2009) or DNA replication (prophage λ) of competing viruses, or when it harbours CRISPR-Cas loci targeting other viruses (Bondy-Denomy & Davidson, 2014). Therefore, proviruses are no longer considered merely as parasites, but are also seen as mutualistic partners of the host (Bondy-Denomy & Davidson, 2014).

*Hfx. volcanii* is a widely used model organism of archaea, with various aspects of its biology extensively studied, including replication, cell division, protein turnover, transcription, translation, and CRISPR-Cas system (Hartman *et al*, 2010; Leigh *et al*, 2011; Maier *et al*, 2019; Perez-Arnaiz *et al*, 2020; Schulze *et al*, 2020).

Notably, viruses are remarkably rare for the genus *Haloferax*, in contrast to other haloarchaeal genera. The first reported *Haloferax*-infecting virus was the HF1 virus infecting *Haloferax lucentense* and *Hfx. volcanii* (Nuttall & Dyall-Smith, 1993), however this virus is no longer available (M. Dyall-Smith, pers. communication). Recently, a new siphovirus, Haloferax tailed virus 1 (HFTV1), was isolated, which infects *Haloferax gibbonsii* LR2-5 (Mizuno *et al*, 2019; Tittes *et al*, 2021a). Another recent addition is *Hfx. volcanii* pleomorphic virus 1 (HFPV-1). This virus exhibits a persistent/chronic infection and is currently the sole available virus known to infect the model haloarchaeon *Hfx. volcanii* (Alarcón-Schumacher *et al*, 2022).

Four proviruses have been identified in *Hfx. volcanii* (Alarcón-Schumacher *et al*., 2022; Dyall-Smith *et al*., 2021). All are encoded on the main chromosome. They are named Halfvol1, Halfvol2, Halfvol3 and Halfvol4 (Table 1 and Suppl. Table 1). Proviruses Halfvol1 and Halfvol2 were shown to be active as they have been identified as circular genomes in a virus stock from *Hfx. volcanii* (Dyall-Smith *et al*., 2021). Halfvol1 and Halfvol3 are distantly related and share characteristics which are typical for pleolipoviruses (Dyall-Smith *et al*., 2021; Liu *et al*, 2015; Roine *et al*, 2010). In general terms, pleolipoviruses are membrane vesicles with spike proteins which encapsulate DNA genomes (Atanasova *et al*, 2015a; Atanasova *et al*, 2015b; Demina & Oksanen, 2020). Halfvol2 shows only limited similarity to known viruses and thus may represent a novel virus group (Dyall-Smith *et al*., 2021). Halfvol4 was described as a defective provirus (Hartman *et al*., 2010; Norais *et al*, 2007).

**Table 1.** Proviruses of *Hfx. volcanii*. This table lists the key characteristics of the four proviruses which have been reported for *Hfx. volcanii*. A complete list of their characteristics is shown in Suppl. Table 1. All four proviruses contain direct repeats of 14 nt. For Halfvol1-3, only the *attP* is contained, thus excluding the repeat copy from *attB*. Thus, the provided length represents the circularised (activated) form of the provirus. The morphotype “novel group” is from (Dyall-Smith *et al*, 2021).

| name | size (bp) | morphotype | flanks (*attP*) | number of integrases |
| --- | --- | --- | --- | --- |
| Halfvol1 | 20,573 | betapleolipovirus | tRNA<sup>Pro</sup>-TGG | 1 |
| Halfvol2 | 12,275 | novel group | tRNA<sup>Ala</sup>-CGC | 2 |
| Halfvol3 | 12,527 | alphapleolipovirus | tRNA<sup>Arg</sup>-TCG | 1 |
| Halfvol4 | 53,702 | not known | - | 3 |

In order to further understand the importance of the proviruses for the host organism *Hfx. volcanii*, we generated a strain that lacks all four proviruses from Halfvol1 to Halfvol4 (ΔHalfvol1-4) and characterised this provirus-free strain in detail.

RNA-seq analyses revealed that ΔHalfvol1-4 shows an increased expression of genes for archaella, chemotaxis and pili and a down-regulation of CRISPR arrays. Furthermore, excised proviral DNA was detected in the culture supernatant for several proviruses and structures identified by transmission electron microscopy (TEM) might represent virus particles, further supporting that proviruses in *Hfx. volcanii* are active viruses.

## Results

### Proviruses of *Hfx. Volcanii*

The genome of the *Hfx. volcanii* type strain DS2 contains four predicted proviruses, and their key characteristics are given in Table 1 and Suppl. Table 1. The genomic organisation of Halfvol1, Halfvol2 and Halfvol3 is illustrated in Suppl. Fig. 1. Halfvol1, Halfvol2, and Halfvol3 are predicted to be double-stranded DNA (dsDNA) viruses, each of them flanked by a perfect direct repeat (14 nt long), or *att* site. This repeat is tRNA-derived (all 14 nt for Halfvol2 and 13 of the 14 nt for Halfvol1 and Halfvol3). At one end is a complete tRNA gene and at the opposite end is a partial copy of that tRNA.

As was reported previously (Dyall-Smith *et al*., 2021; Liu *et al*., 2015; Roine *et al*., 2010), Halfvol1 and Halfvol3 encode proteins which are homologous to those from pleolipoviruses (Table 2). In particular, the structural proteins VP4 (spike protein) and VP3 (matrix protein) are present in Halfvol1 (HVO_0271; HVO_0269) and in Halfvol3 (HVO_1431; HVO_1432). Moreover, genes homologous to VP8 (ATPase, HVO_0274; HVO_1426) are present in both proviruses. In addition, both proviruses encode winged-HTH domain proteins (HVO_0260 and HVO_0261; HVO_1423) with distant similarity to the halovirus phiH repressor protein which is important for lysogeny (Ken & Hackett, 1991).

**Table 2.**
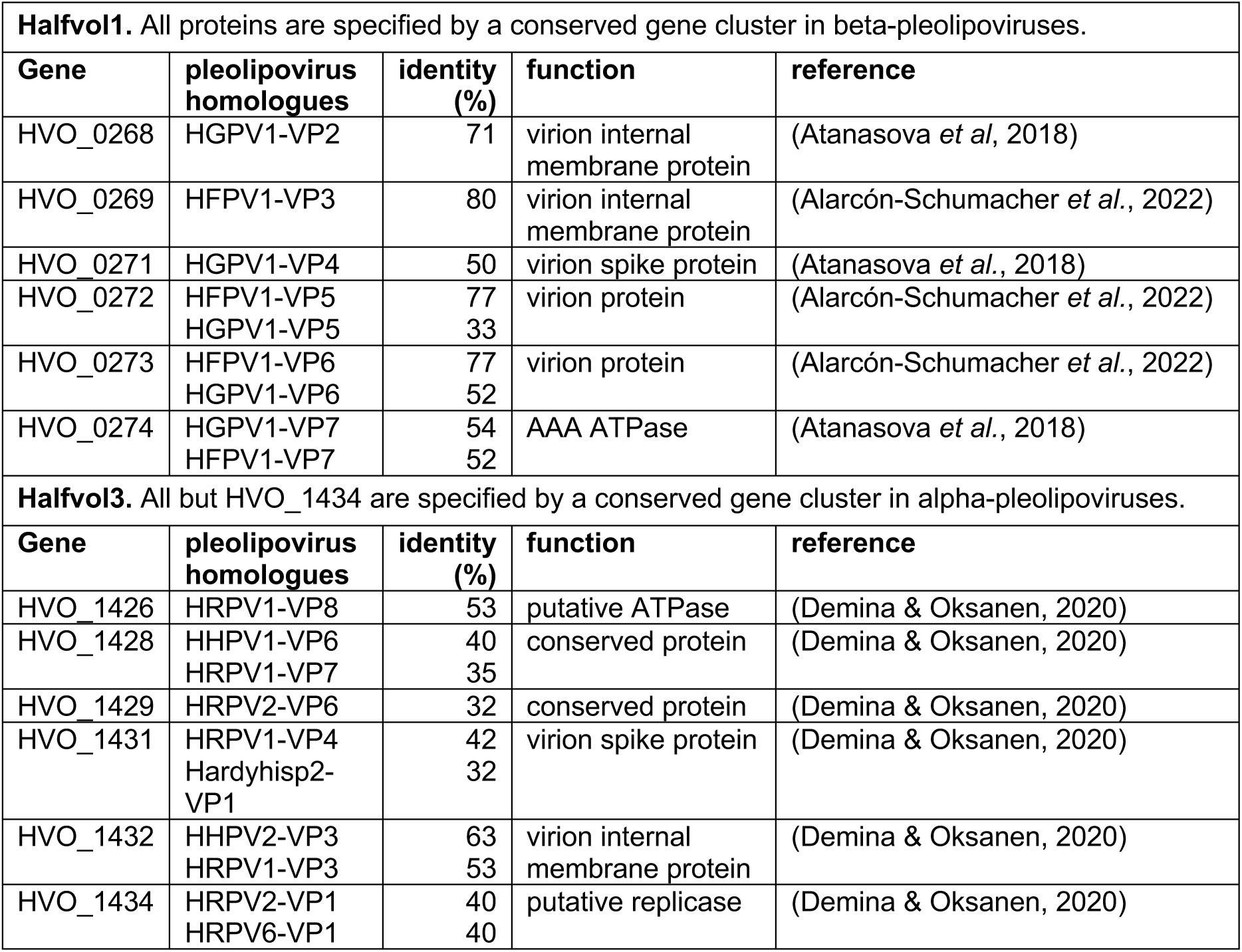
Proteins encoded on proviruses Halfvol1 and Halfvol3 which are related to key virus proteins. Proteins which are encoded on proviruses Halfvol1 and Halfvol3 and which are homologous to viral proteins are shown. Excluded are (a) integrases and (b) HTH domain proteins distantly related to the halovirus phiH repressor. No proteins homologous to virus proteins are encoded on Halfvol2 and Halfvol4. Virion protein indicates proteins present in purified virus particles. Spike protein is the viral attachment protein.

Attempts to identify a Halfvol2 protein related to a virus capsid protein were not successful. Sequence similarity searches did not retrieve a viral capsid protein. Also, a foldseek analysis did not reveal any capsid protein showing significant (better than e-3) structural similarity. However, a packaging ATPase candidate could be retrieved. HVO_0377 shows structural similarity to the likely packaging ATPases from *Haloarcula hispanica* virus PH1 (e-10; HhPH1_gp13; UniProt:M4JFA3) and from *Sulfolobus* turreted icosahedral virus 2 (e-11; STIV2_B204; UniProt:D5IEZ9; PDB:4KFU).

Halfvol4 is characterised by high A/T content when compared to the main chromosome and it displays an unusual codon usage variation, when compared to the codon usage of the whole chromosome (Hartman et al., 2010). Halfvol4 contains three XerC/D-like integrases, a type IV secretion system (T4SS), two divergent *orc* genes, a restriction modification system (Hartman et al., 2010) and the recently identified new defence system Avast (HVO_2276, Avast type 2) (Gao *et al*, 2020; Payne *et al*, 2022). The Halfvol4 region was previously reported as a defective provirus (Hartman *et al*., 2010; Norais *et al*., 2007). Initially, this provirus was attributed to genome positions 2110297-2163576 (covering HVO_2252 to HVO_2293A) (Hartman et al., 2010) or covering HVO_2252 to HVO_2293 (Norais *et al*., 2007).

We shifted the boundaries of Halfvol4 to genome positions 2109865-2163566 (covering HVO_2251A to HVO_2293A), after finding that this element is enclosed by a 14 nt perfect direct repeat. A 3^rd^ perfect copy and a near-complete copy (13 nt) of this repeat is located internally within Halfvol4. In some closely related haloarchaeal species, this region shows heterogeneity (Suppl. Table 2). While in *Hfx. volcanii* Halfvol4 is inserted at this site, in other haloarchaeal species there is no insertion present, while still others carry an insert, but of different size (Suppl. Table 2). The genomic organisation of Halfvol4 is illustrated in Suppl. Fig. 1.

### Generation of the provirus-free *Hfx. volcanii* strain ΔHalfvol1-4

To investigate the roles and impacts of proviruses in *Hfx. volcanii*, we generated a deletion strain in which all four predicted proviruses were removed (ΔHalfvol1-4). In a previous study, the provirus Halfvol3 was deleted from strain H133 (Δ*pyrE2* Δ*leuB* Δ*hdrB* Δ*trpA*) and substituted with the *trpA* marker (Alarcón-Schumacher *et al*., 2022). Subsequently, the Halfvol4 locus was removed from the ΔHalfvol3 strain, followed by the deletion of Halfvol1 from the ΔHalfvol3ΔHalfvol4 strain. The deletion of Halfvol1 turned out to be quite difficult, possibly because the mini-chromosome pHV4 is integrated into Halfvol1 (see below), thus deletion of Halfvol1 also removes pHV4 from the genome.

Since pHV4 carries multiple essential genes (Norais *et al*., 2007) Halfvol1 can only be deleted if pHV4 becomes episomal again. Lastly, Halfvol2 was deleted from the ΔHalfvol1ΔHalfvol3ΔHalfvol4 strain resulting in the provirus free strain (ΔHalfvol1-4), which was confirmed by Southern analysis (Suppl. Fig. 2). We use the strain H100 (Δ*pyrE2* Δ*leuB* Δ*hdrB*) as a control which differs from the parent strain H133 by carrying the *trpA* gene under its natural promoter. This is consistent with the *trpA* gene being present in ΔHalfvol1-4 at the position of Halfvol3, but under the p.*fdx* promoter.

Previous findings indicate that, in the *Hfx. volcanii* DS70 strain, the pHV4 mini-chromosome was integrated into the main chromosome through recombination between the ISHvo9 transposons coding for transposase HVO_0278 (chromosome) and HVO_A0279 (pHV4) (Hawkins *et al*, 2013). Given that HVO_0278 is encoded within the Halfvol1 locus, this suggests that pHV4 is integrated into Halfvol1. Pulsed Field Gel Electrophoresis (Suppl. Fig. 5) revealed the excision of mini-chromosome pHV4 from the chromosome in ΔHalfvol1-4.

Since the Halfvol1 locus, and therefore the site of the integration of pHV4, has been deleted in this strain, pHV4 exists exclusively in its episomal form in the provirus-free strain. These analyses revealed that pHV4 is present in the episomal form in the ΔHalfvol1-4 strain, whereas in the other strains it is present both in the chromosome-integrated form and additionally as an episomal mini-chromosome.

### Phenotypic characterisation of the provirus-free strain ΔHalfvol1-4

A number of phenotypic tests were used to compare the ΔHalfvol1-4 strain with the isogenic H100 strain. Growth analysis at optimal temperature and salt concentration (45°C and 18% BSW) indicates that the ΔHalfvol1-4 strain has a slightly better growth rate than the control strain (growth rate H100: 0.07, Δ1-4: 0.09) (Fig. 1). This suggests that the presence of proviruses is associated with growth retardation. Growth curves were also tested under salt stress (15% and 23% BSW) and temperature stress (30°C and 50°C). Under low salt conditions and both low and high temperatures, the provirus-free strain is also growing better than the control (Suppl. Fig. 2), consistent with the results under standard conditions. However, at higher salt conditions (23% BSW), the ΔHalfvol1-4 strain exhibited a slight growth retardation. Phase-contrast microscopy analysis revealed that the ΔHalfvol1-4 strain predominantly exhibited an elongated cell shape, in contrast to the control H100 strain, which displayed a round cell shape. However, round-shaped cells were also observed in ΔHalfvol1-4, primarily during the late exponential phase (OD_650_: 0.6) (Fig. 2). Viruses and proviruses can significantly influence the motility of their host organisms (Brathwaite *et al*, 2015; Wang *et al*, 2009). To explore this, a swarming assay was conducted with ΔHalfvol1-4 and a control strain (H100). The ΔHalfvol1-4 strain exhibited hypermotility compared to the control, suggesting that the presence of proviruses reduces motility (Fig. 3). In addition, biofilm formation was investigated but no significant difference was observed between the provirus-free strain and the control. (Suppl. Fig. 4). The effects of provirus deletion on cell responses to stressors were investigated in two experiments: oxidative stress and UV survival assays. The ΔHalfvol1-4 strain demonstrated high resilience to oxidative stress at a high H_2_O_2_ concentration (6 mM), compared to the control (Fig. 4). However, no significant differences were observed in response to UV stress (Suppl. Fig. 3).

**Figure 1.**
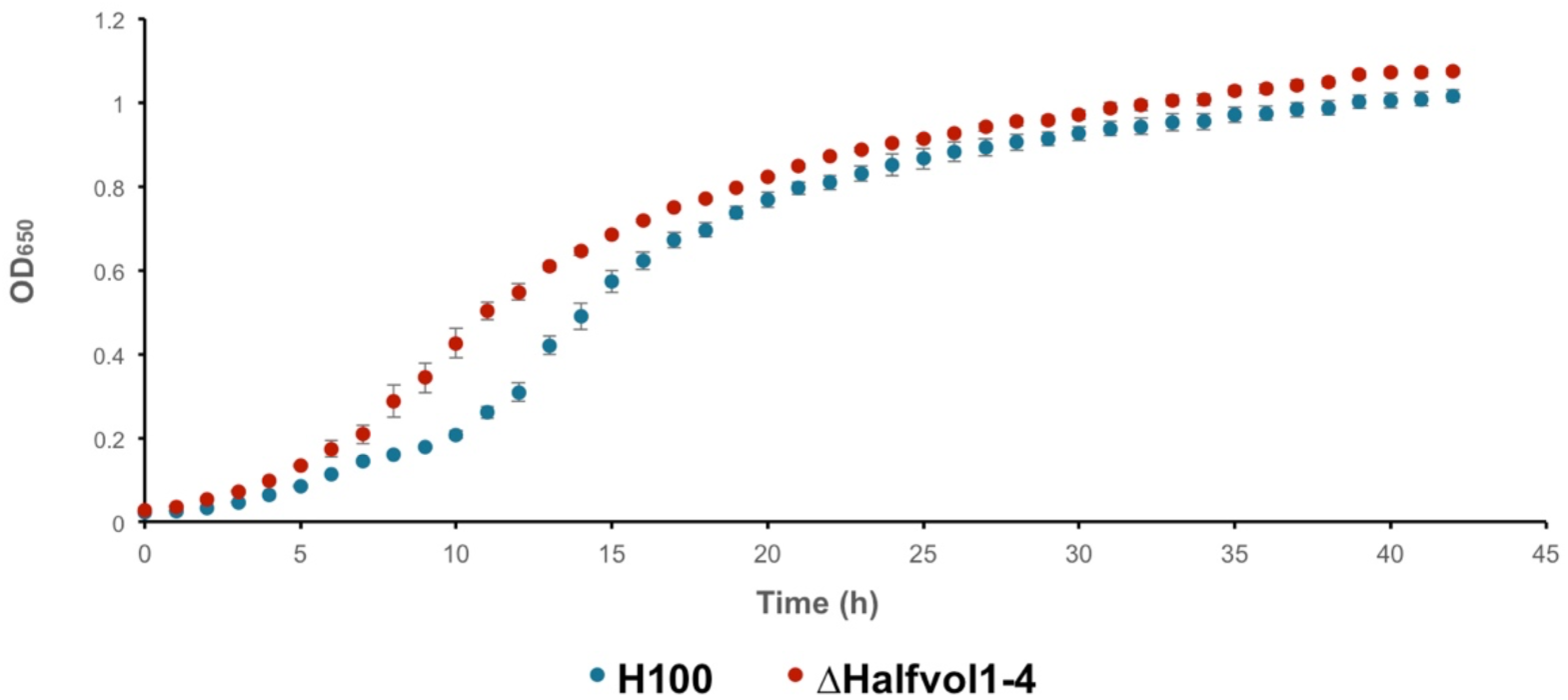
Growth curves of provirus-free strain and control strain. Both strains were grown under standard conditions (45°C and 18%BSW). The provirus containing control is *Hfx. volcanii* strain H100 (red) and its growth has been compared with that of the provirus-free strain ΔHalfvol1-4 (blue). Data shown are based on three biological replicates each. See methods for details.

**Figure 2.**
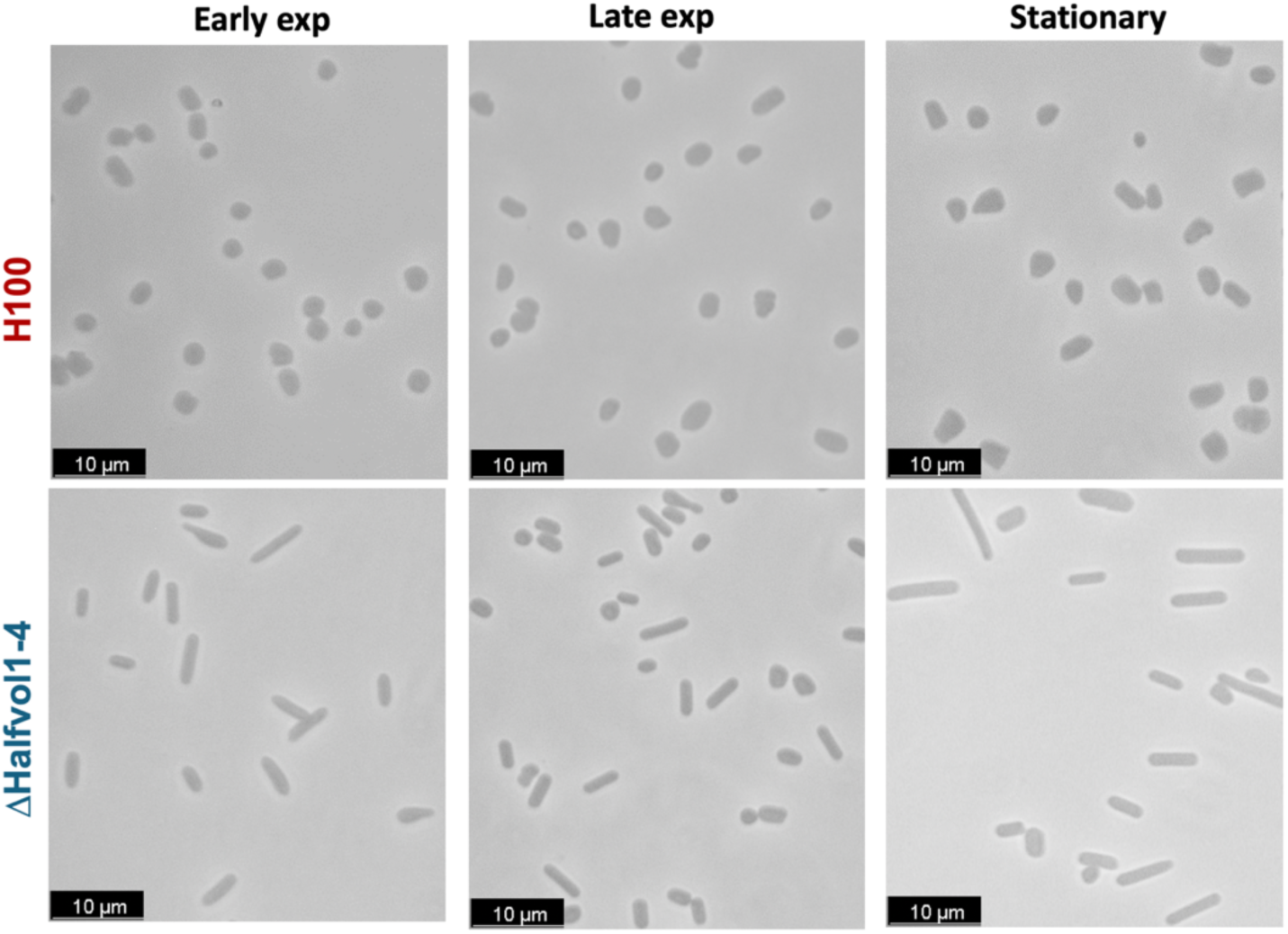
Microscopy analysis of provirus-free strain and control strain. Phase contrast microscopy images of the control strain H100 and the provirus-free strain ΔHalfvol1-4, three biological replicates were analysed for each strain. The first column shows images of H100 and ΔHalfvol1-4 in early exponential phase (OD_650_ 0.3), the second column in late exponential phase (OD_650_ 0.6) and the third column in stationary phase (OD_650_ 1.0).

**Figure 3.**
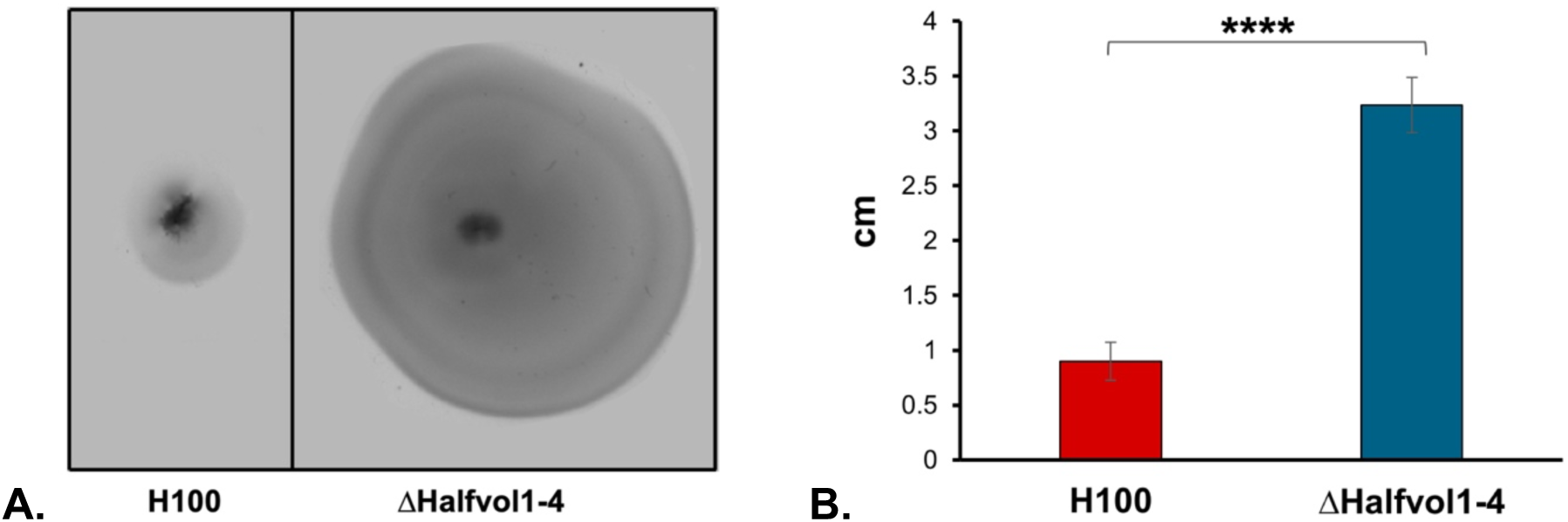
Swarming assay of provirus-free strain and control strain. Swarming motility of the control strain H100 and the provirus-free strain ΔHalfvol1-4 was tested at 45°C in Hv-Ca plates (0.3% agar). (A) Phenotypical observation of the halo disk due to the swarming of strains H100 and ΔHalfvol1-4. (B) The measured diameter of the halo disks. Asterisks indicate significant differences (t-test); ****: highly significant (p-value ≤ 0.0001).

**Figure 4.**
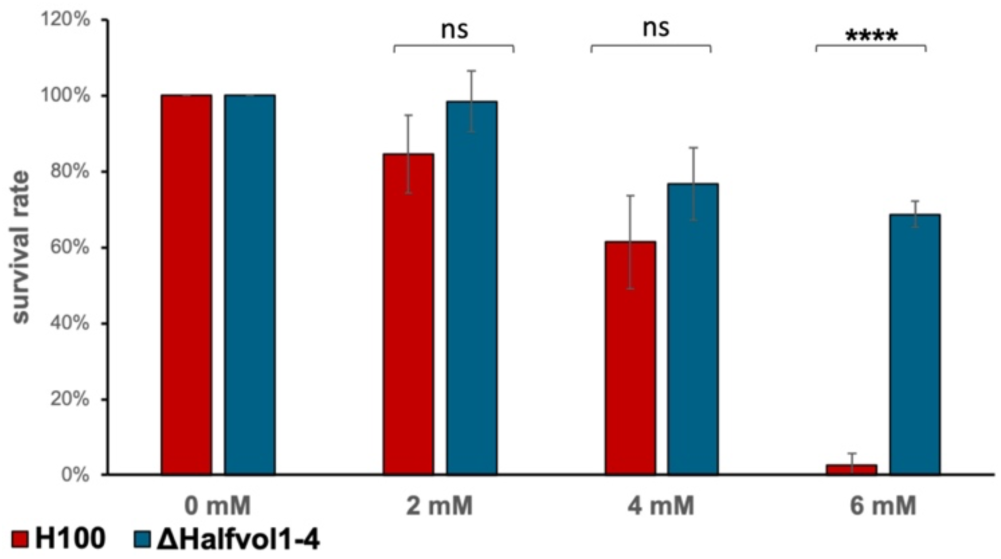
Oxidative stress resilience between provirus-free strain and control strain. The cultures were incubated for 1 hour in 18% salt water with different concentrations of H_2_O_2_ (0 mM, 2 mM, 4 mM and 6 mM). Cells were then plated on Hv-YPC + thymidine agar plates and the survival rate of the strains was determined. The values shown are based on three biological replicates. Symbols shown on top of the bar: asterisks indicate significant differences (t-test); ****: highly significant (p-value ≤ 0.0001), ns= not significant.

### Transcriptome analysis

To assess whether provirus removal affects gene expression, the transcriptome of the provirus deletion strain ΔHalfvol1-4 was compared to that of the control strain H100 by RNA-seq (Table 3, 4 and Suppl. Table 3). The RNA-seq data indicates an increased expression of archaella, chemotaxis and pili genes in the ΔHalfvol1-4 strain, which are involved in motility. Specifically, the locus from HVO_1203 to HVO_1225, which contains genes associated with motility, shows upregulation of approximately 2- to 3-fold in ΔHalfvol1-4. This genetic expression pattern is consistent with the hypermotility observed in the swarming assay (Fig. 3), thereby providing a molecular explanation for the observed phenotypic behaviour. Moreover, in ΔHalfvol1-4, a pronounced down-regulation of CRISPR arrays (P1, P2, and locus C) was observed, reaching up to a 9-fold reduction compared to the control strain. The reduction in crRNA expression in the provirus-free strain suggests a potential influence of proviruses on the adaptive immune system of *Hfx. volcanii*. Additionally, transcription regulators (HVO_A0394, HVO_A0135, HVO_C0076, HVO_A0527, HVO_1052) and genes involved in the amino acid metabolism (HVO_A0295, HVO_0285) also exhibited decreased expression levels (Suppl. Table 3).

**Table 3.** Differential gene expression between ΔHalfvol1-4 and strain H100 (control). Many genes are differentially regulated in the provirus deletion strain and in the table only the 10 most strongly down- and up-regulated genes are shown here. Down-regulation is indicated by negative log2FC values, up-regulation by positive log2FC values. The log2 fold change (column log2FC) deletion vs. wild type is given alongside the HVO gene number (column Gene), gene annotation (column Annotation) and adjusted p value (column p adj).

| TOP 10 up-regulated genes |  |  |  |
| --- | --- | --- | --- |
| Gene | Annotation | log2FC | p adj |
| HVO_0281 | tRNA-Pro | 3.08 | 2.5E-109 |
| HVO_0288 | tRNA-His | 2.10 | 1.84E-50 |
| HVO_0287 | conserved hypothetical protein | 2.08 | 2.81E-32 |
| HVO_0283 | archaea-specific helicase AshA | 2.06 | 9.14E-53 |
| HVO_0282 | DASS family transport protein | 2.05 | 6.19E-25 |
| HVO_B0065 | thioredoxin domain protein | 1.75 | 1.02E-08 |
| HVO_B0053 | DUF3209 family protein | 1.63 | 1.24E-07 |
| HVO_0284 | cupin 2 barrel domain protein | 1.60 | 4.14E-12 |
| HVO_0285 | M50 family metalloprotease | 1.60 | 7.41E-13 |
| HVO_1228 | DUF5059 domain / halocyanin domain protein, hcpE | 1.55 | 1.66E-07 |
| TOP 10 down-regulated genes |  |  |  |
| Gene | Annotation | log2FC | p adj |
| HVO_2293_T | H/ACA guide RNA (snoRNA) | -7.10 | 1.2E-122 |
| HVO_2293A | small CPxCG-related zinc finger protein | -5.69 | 2.95E-37 |
| HVO_2293s | small RNA | -5.66 | 7.15E-41 |
| HVO_A0295 | amidase (hydantoinase/carbamoylase family, amaB2) | -3.53 | 6.35E-21 |
| HVO_A0129s | small RNA | -3.34 | 1.28E-33 |
| CRISPR2 | CRISPR Locus P1 | -3.18 | 3.33E-43 |
| HVO_A0394 | HTH domain protein | -3.13 | 4.32E-77 |
| HVO_1252 | conserved hypothetical protein | -3.11 | 3.43E-96 |
| HVO_A0212s | small RNA | -3.07 | 1.9E-31 |
| CRISPR3 | CRISPR Locus P2 | -2.93 | 7.79E-31 |

**Table 4.**
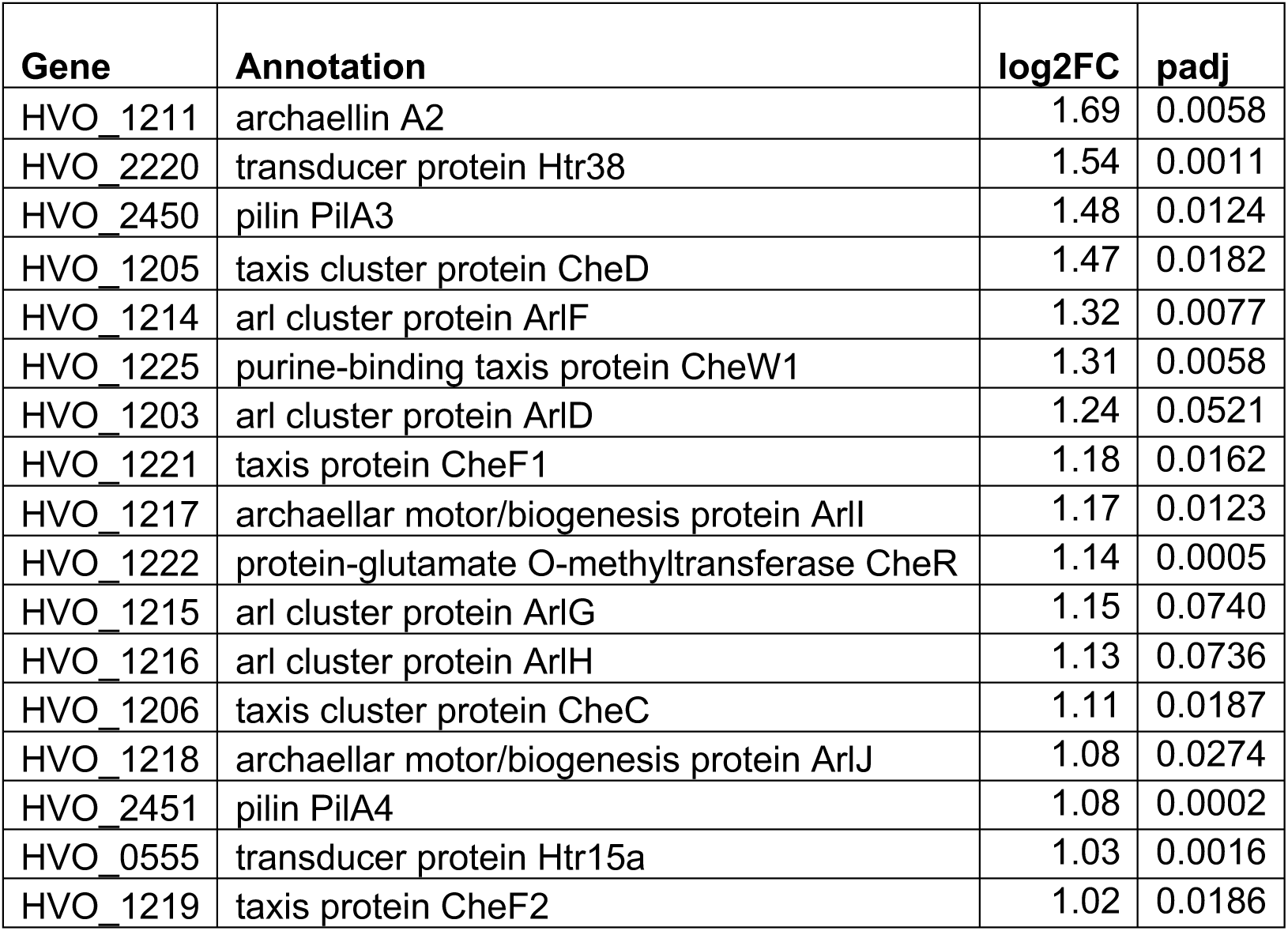
Differential gene expression of motility genes. Motility related genes showing up-regulation are listed. All genes regulated with a log2 fold change (log2FC) > 1 and an adjusted p-value (padj) < 0.074 are shown.

However, it should be noted that proteins with the code prefix HVO_A originate from mini-chromosome pHV4 which is integrated into the major chromosome in the parent strain but occurs in episomal form in strain ΔHalfvol1-4. Changes in expression of these genes might also be related to this. Curiously, several genes immediately downstream of Halfvol1 (HVO_0281 to HVO_0289) were up-regulated up to 8-fold (log2FC 3.1) in the ΔHalfvol1-4 deletion strain. This suggests that the deletion of Halfvol1 may lead to the upregulation of these genes, indicating a possible regulatory effect of Halfvol1 on its downstream region.

### Proviruses excise from the chromosome and are released from cells

Two different approaches were used to detect provirus induction: (1) PCR and Sanger sequencing were used to analyse the *attP* and *attB* sites, to identify configurations indicative of provirus excision from the chromosome (in strains H100, H133 and Δ*oapA*, see Fig. 5, 7 and 8B.). (2) Examination of culture supernatants by (a) electron microscopy to detect virus-like particles (Fig. 9 and Suppl. Fig. 7, strain Δ*oapA*), and (b) mass spectrometry to detect viral proteins.

In order to demonstrate provirus excision and release from the cell, isolation of viruses from the culture supernatant was performed. Proviruses Halfvol1, Halfvol2, and Halfvol3 are flanked by 14 nt direct repeats, and circularise upon excision. Virus circularisation was validated using PCR with outward-facing primers positioned close to the provirus ends (Fig. 5). PCR products that, traverse the *attP* site were confirmed by Sanger sequencing. Intact *attP* sites indicate a circularised virus genome, thus confirming virus excision. Interestingly, provirus excision was more efficient when the culture was grown at low temperature (30°C) (Fig. 6). We did not detect any circularisation of Halfvol4. Given the presence of circular virus DNA, the occurrence of provirus-free *attB* sites was also examined via PCR in three provirus-containing strains which were used in this study: the control strain H100, the parent strain H133, and in addition strain Δ*oapA*, which has a deletion in gene *oapA* (HVO_3014) and produces only a small amount of extracellular vesicles (see below) (Mills et al, 2024). Convergent primers targeting the upstream and downstream region of the provirus were used. Given the substantial size of the proviral regions (12-20kb), a PCR product would only be expected if the provirus had been naturally excised from the chromosome (Fig. 7). PCR products of the anticipated size were obtained and subjected to Sanger sequencing which confirmed the existence of provirus-free *attB*, and so validated that the proviruses had been excised from the chromosome.

**Figure 5.**
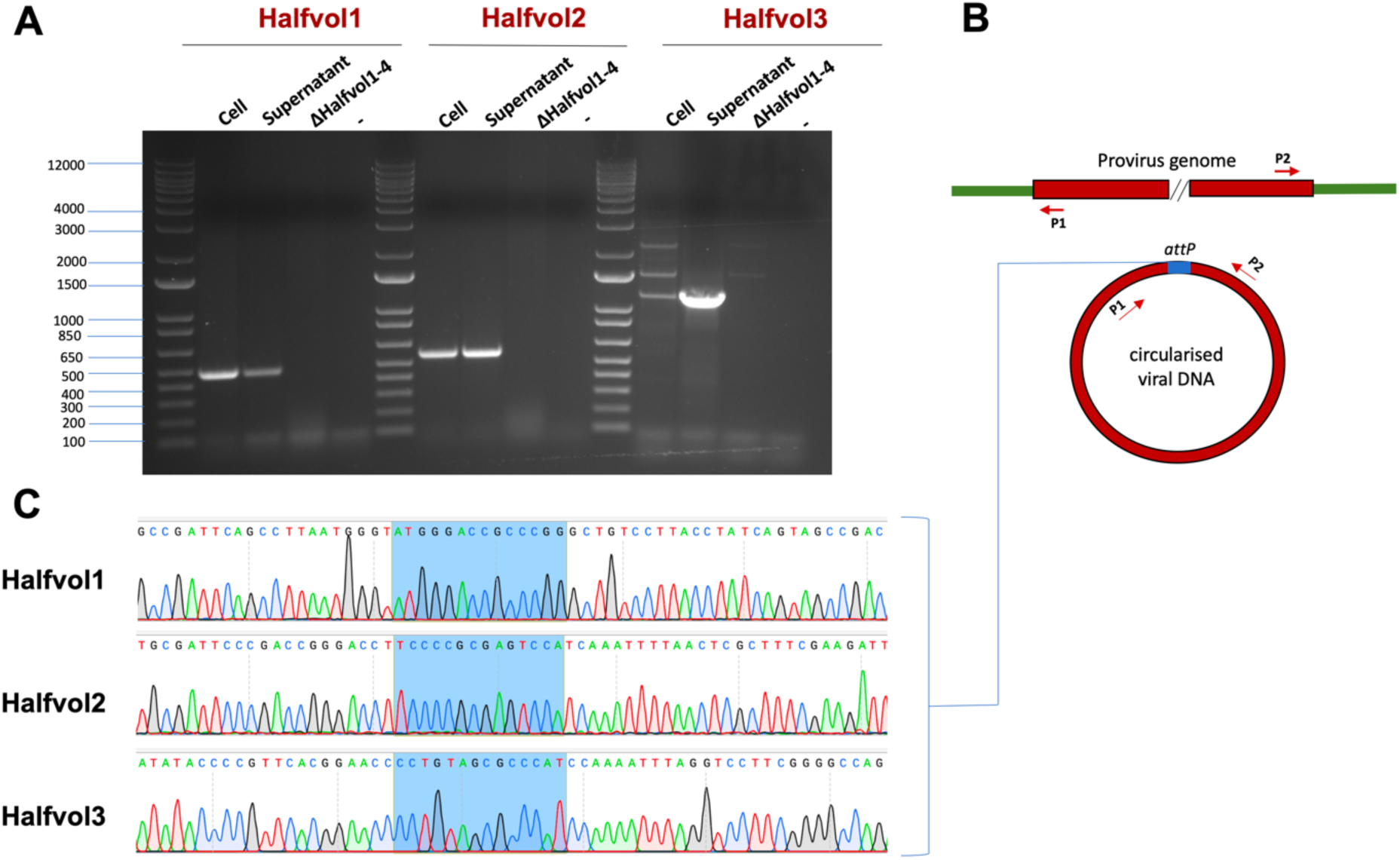
Detection of *attP* site of Halfvol1, Halfvol2 and Halfvol3, indicating provirus excision. **A.** PCR products obtained with outward facing primers, indicating the circularisation of Halfvol1, Halfvol2, and Halfvol3, lane “Cell”: wild type cells (H119) were taken from a plate and used as template. In “Supernatant”, 1µl of 1:10 dilution of isolated excised proviruses from the culture supernatant from liquid cultures was used as template. In “ΔHalfvol1-4” ΔHalfvol1-4 cells taken from a plate were used as template, while “-” is the negative control. **B.** The position of the primers employed in the PCR are indicated by red arrows. A PCR product is generated exclusively when the viral DNA has circularised. **C.** Sanger sequencing of the bands obtained in the PCR from the “Supernatant” samples. The *attP* sequence is highlighted in light blue.

**Figure 6.**
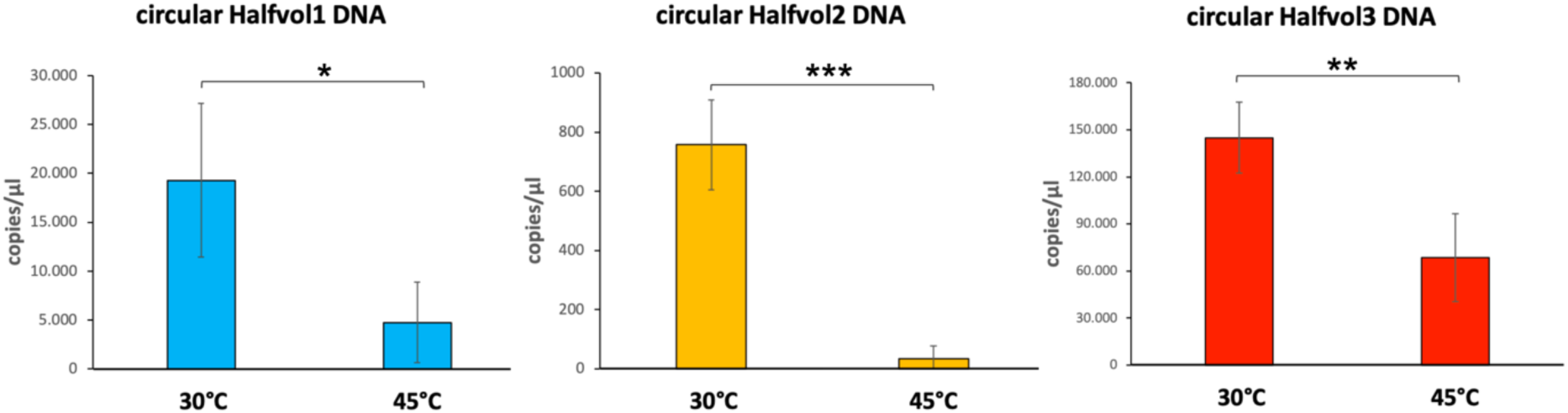
Quantification of circular Halfvol1, Halfvol2 and Halfvol3 DNA. qPCR was employed to determine the concentration of circular Halfvol1, Halfvol2 and Halfvol3 DNA in the culture supernatant of the provirus-containing strain Δ*oapA*, cultured at 30°C (low growth temperature for *Hfx. volcanii*) or 45°C (standard growth temperature) during the stationary phase (OD_650_ 0.86). Results have been taken from three biological replicates. Asterisks indicate significant differences (t-test); * : P ≤ 0.05; ** : P ≤ 0.01; *** : P ≤ 0.0001.

**Figure 7.**
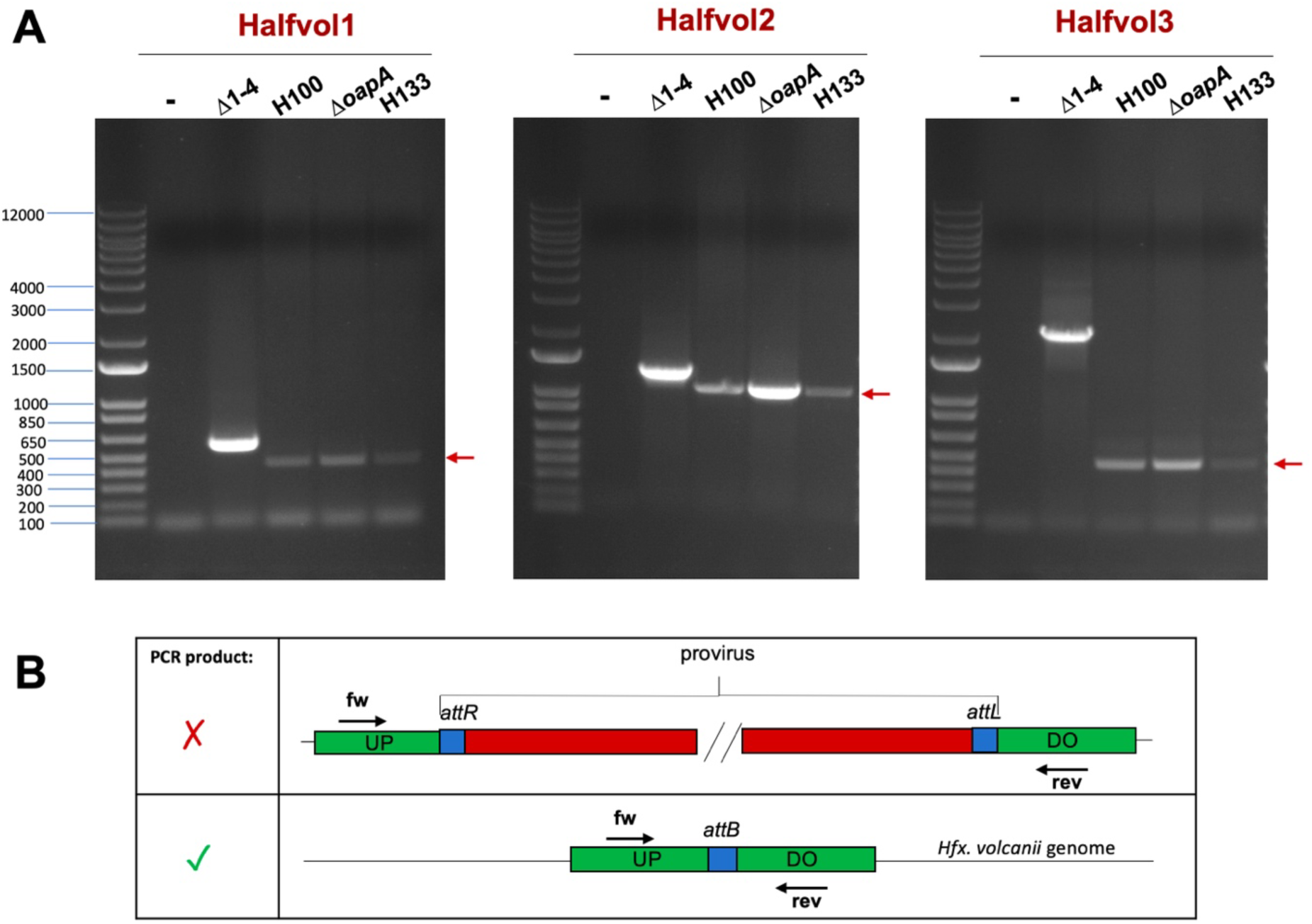
Detection of *attB* sites, indicating provirus excision. **A.** The excision of Halfvol1, Halfvol2 and Halfvol3 from the chromosome is shown in different *Hfx. volcanii* strains (Δ1-4, H100, Δ*oapA* and H133) by PCR. For Halfvol1, spontaneous excision results in a 451 bp signal (weak signals in lanes H100, ΔoapA, H133, indicated by an arrow) whereas the deletion strain (ΔHalfvol1-4: Δ1-4) shows a band at 585 bp. For Halfvol2, spontaneous excision gives a signal of 1,015 bp (indicated by an arrow), whereas the deletion strain (Δ1-4) gives a signal of 1263 bp. For Halfvol3, spontaneous excision results in a 465 bp signal (indicated by an arrow), while the deletion strain (Δ1-4) shows a band at 2,172 bp. The size increase is due to insertion of the *trpA* marker**. B.** The primers employed in the PCR are indicated by black arrows and they are located in the upstream (UP) and downstream (DO) regions of the specific provirus. A PCR product is generated exclusively when the provirus excised from the *Hfx. volcanii* chromosome.

### Identification of potential virus particles

Virus particles can be enriched by subjecting the culture supernatant to CsCl density gradient centrifugation. Due to the production of extracellular vesicles (EVs), *Hfx. volcanii* naturally shows a strong pink band in such gradients (Mills *et al*., 2024). Since EVs interfere with virus detection, we used strain *ΔoapA*, which has a largely reduced EV production by deletion of the gene coding for GTPase HVO_3014 (*oapA*; also called *arvA*) (Alarcón-Schumacher *et al*., 2022). This strain was used to enrich virus particles upon CsCl density gradient centrifugation and to investigate the gradient fraction for the presence of virus particles using electron microscopy.

After CsCl density gradient centrifugation of the virus preparation from *ΔoapA*, two distinct whitish bands were visible (Fig. 8A., right tube). Whereas the ΔHalfvol1-4 preparation, which produces EVs, showed a singular wide pink band (Fig. 8A, left tube). The presence of viral DNA (circular Halfvol1, Halfvol2 and Halfvol3) was confirmed by PCR in both bands of the *ΔoapA* sample (Fig. 8B). Samples from the two bands in *ΔoapA,* and the corresponding parts in ΔHalfvol1-4 as control, were subjected to transmission electron microscopy analysis. Within the first band, mainly two different particles were observed (Fig. 9A). Unstructured round particles (diameter of approximately 40 nm) were observed in *ΔoapA* and ΔHalfvol1-4, which probably represent EVs. In *ΔoapA,* larger and more structured particles (diameter of approximately 80 nm) were detected. These latter particles might represent virus particles originating from activated proviruses. A set of representative images is shown in Suppl. Fig. 7. In the second band of the *ΔoapA* sample, particles that resemble head-tailed structures were observed (Fig. 9B). The heads of the particles are spherical and uniform in size, characterised by a bright white core surrounded by a darker halo. The “tails” appeared to be structured, with a notable variability in length, with the longest tail measured to be approximately 1.7 µm. It is yet unresolved if these are virus tails or if they are cellular structures (archaella or pili) to which virus particles have attached. A series of representative images is shown in Suppl. Fig. 7.

**Figure 8.**
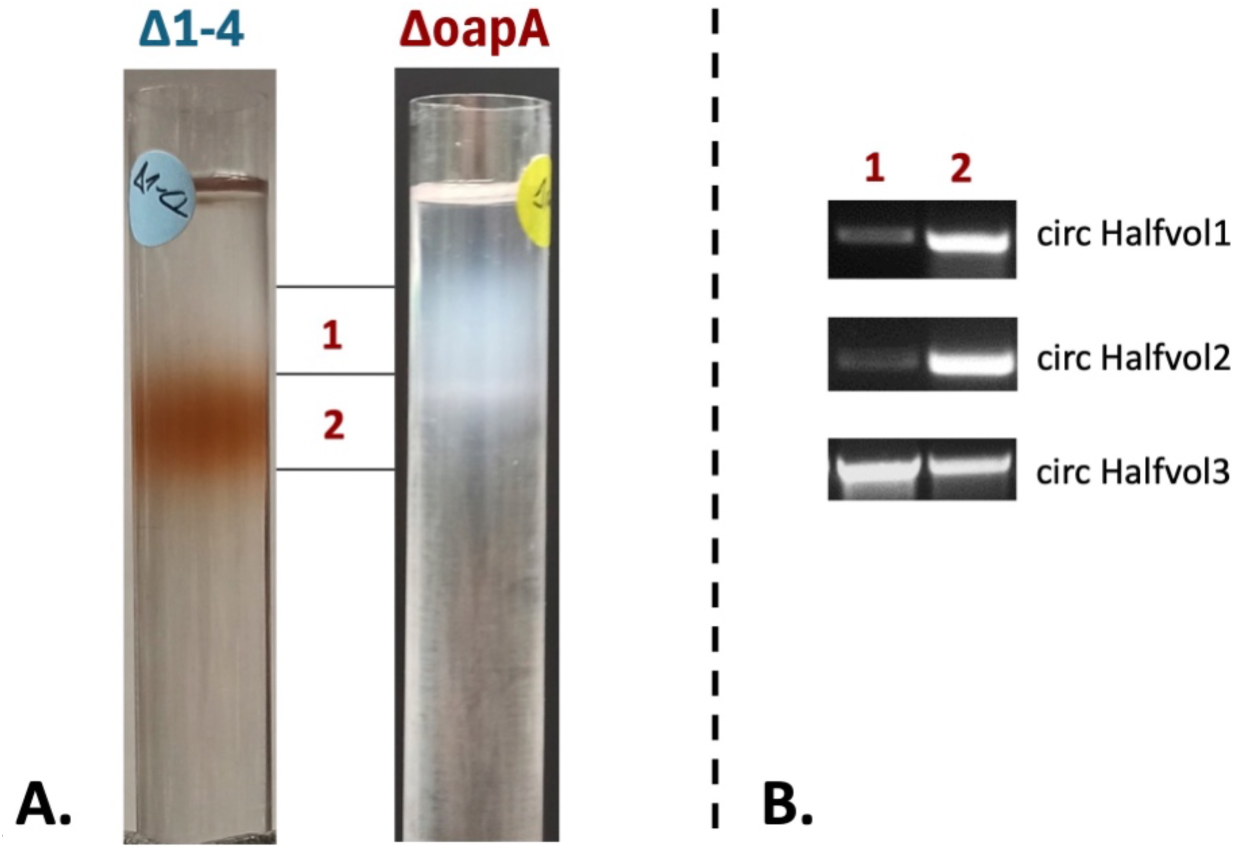
Purification of isolated excised provirus from Δ*oapA* supernatant by CsCl density gradient. **A.** CsCl density gradient ultracentrifugation for samples obtained from strains ΔHalfvol1-4 (Δ1-4, lacking proviruses) and Δ*oapA* (containing proviruses). The ΔHalfvol1-4 sample (left tube) shows a broad pink band while the Δ*oapA* sample reveals one broad white band and one distinct white band (right tube). **B.** PCR confirms the presence of circularised viral DNA in the first and second bands (“1” and “2”) of the Δ*oapA* sample. Circular forms of Halfvol1, Halfvol2 and Halfvol3 DNA were found in fractions of both bands.

**Figure 9.**
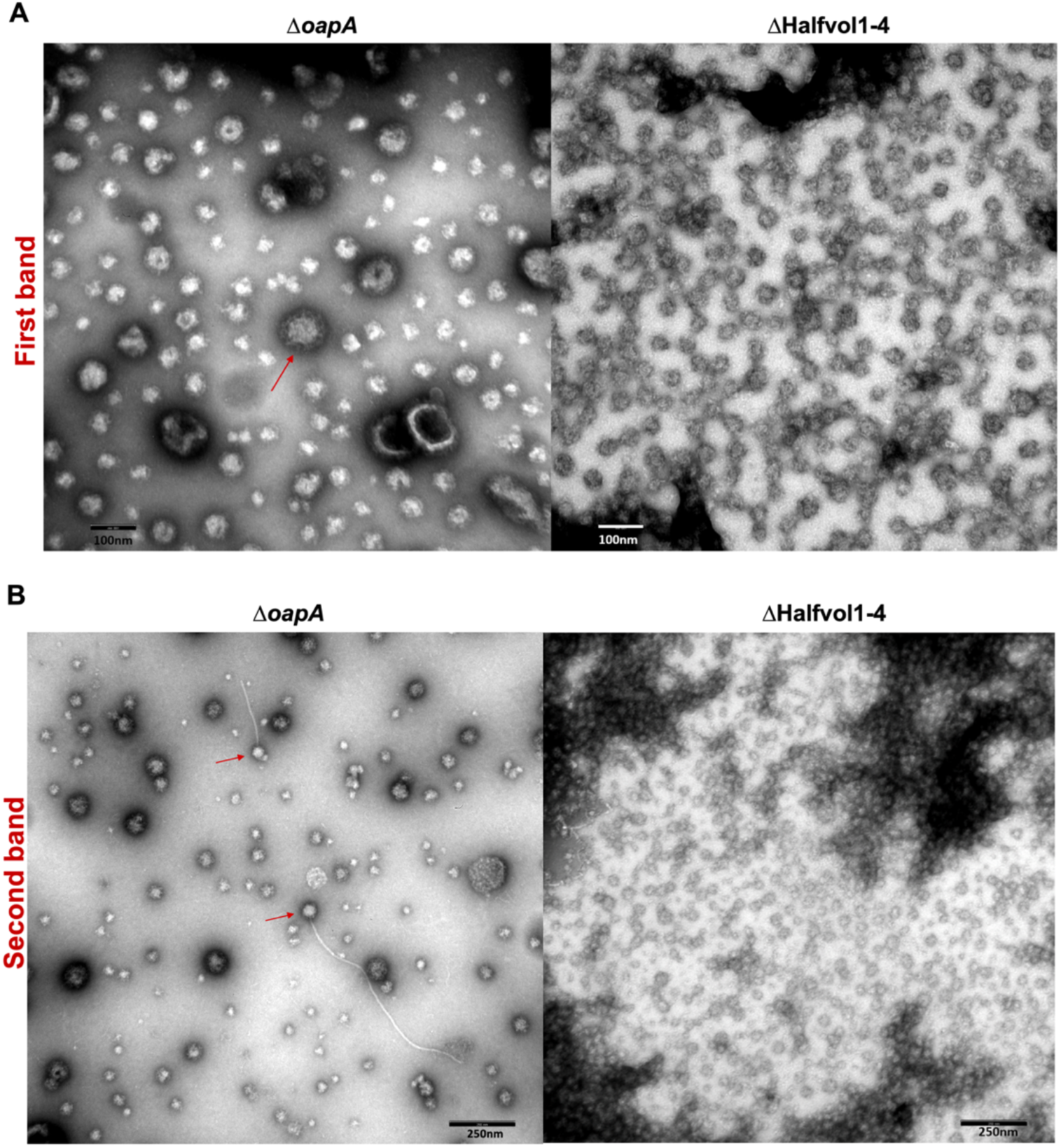
Transmission electron microscopy images of purified supernatant from provirus-containing strain Δ*oapA* and provirus-free strain ΔHalfvol1-4. The isolation of virus sample was performed in the provirus-containing strain (Δ*oapA*) and the same procedure was carried out in the provirus-free strain (ΔHalfvol1-4) as a control. **A.** First band of the CsCl density gradient from strain Δ*oapA* and its counterpart from strain ΔHalfvol1-4. A structure that resembles a pleolipovirus, complete with spike proteins, is highlighted with a red arrow. The scale bar indicates 100 nm. **B.** Second band of the CsCl density gradient from strain Δ*oapA* and its counterpart from strain ΔHalfvol1-4. Head-tailed structures were observed in the Δ*oapA* sample (red arrows). The scale bar indicates 250 nm.

### Analysis of density gradient centrifugation bands by mass spectroscopy

The second band from CsCl density gradient centrifugation derived from strain *ΔoapA* and its counterpart from strain ΔHalfvol1-4 were loaded on a 12% SDS gel and were subjected to mass spectrometry analysis. Viral proteins were identified in the *ΔoapA* sample (Table 5) but not in the ΔHalfvol1-4 sample, that was analysed as a control. Four different proteins associated with Halfvol1 were identified, one of them being HVO_0271, the homolog of structural protein VP4. Seven different proteins associated with Halfvol3 were identified, one of them being a VP4 homologue (HVO_1431) with a total spectrum count of 69 (Table 5). Remarkably, nineteen different proteins were identified for Halfvol4, featuring the highest number of total spectrum counts compared to the other viruses.

**Table 5.** Provirus proteins identified in purified fractions. Fraction 2 from the CsCl gradient from ΔMV and Δ1-4 cells were analysed with mass spectrometry. Identified provirus proteins are listed, for which the total spectrum count was higher in Δ*oapA* samples compared to Δ1-4 samples. An average of the total spectrum count values from the three biological replicates was calculated. Column Δ*oapA*: average total spectrum counts from Δ*oapA* proteins detected, column Δ1-4: average total spectrum counts from Δ1-4 proteins detected.

| gene | annotation | ΔoapA | Δ1-4 |
| --- | --- | --- | --- |
| **Halvol1** |  |  |  |
| HVO_0277 | conserved hypothetical protein | 6 | 0 |
| HVO_0272 | homolog to HGPV1-ORF5 | 5 | 0 |
| **HVO_0271** | **structural protein VP4 (spike protein)** | **3** | **0** |
| HVO_0259 | conserved hypothetical protein | 2 | 0 |
| **Halfvol2** |  |  |  |
| HVO_0379_A | conserved hypothetical protein | 19 | 0 |
| HVO_0379_B | conserved hypothetical protein | 19 | 0 |
| **Halfvol3** |  |  |  |
| **HVO_1431** | **structural protein VP4 (spike protein)** | **69** | **0** |
| HVO_1424 | conserved hypothetical protein | 16 | 0 |
| HVO_1425 | conserved hypothetical protein | 10 | 0 |
| HVO_1429 | homolog to pHK2-ORF6 | 5 | 0 |
| HVO_1428 | homolog to pHK2-ORF7 | 3 | 0 |
| HVO_1433 | homolog to pHK2-ORF2 | 3 | 0 |
| HVO_1426 | homolog to virus protein HRPV1-VP8 | 2 | 0 |
| **Halfvol4** |  |  |  |
| HVO_2286 | conserved hypothetical protein | 208 | 2 |
| HVO_2263 | conserved hypothetical protein | 117 | 0 |
| HVO_2269 | RmeR | 51 | 1 |
| HVO_2267 | conserved hypothetical protein | 29 | 0 |
| HVO_2266 | conserved hypothetical protein | 22 | 0 |
| HVO_2256 | DUF262 family protein | 17 | 0 |
| HVO_2271 | RmeS | 16 | 0 |
| HVO_2279 | conserved hypothetical protein | 15 | 0 |
| HVO_2270 | RmeM | 14 | 0 |
| HVO_2276 | Avs2 family antiviral STAND domain protein | 14 | 0 |
| HVO_2265 | conserved hypothetical protein | 11 | 0 |
| HVO_2253 | conserved hypothetical protein | 10 | 0 |
| HVO_2288 | hypothetical protein | 8 | 0 |
| HVO_2291 | hypothetical protein | 6 | 0 |
| HVO_2255 | conserved hypothetical protein | 5 | 0 |
| HVO_2275 | N-6 adenine-specific DNA methylase domain protein | 4 | 3 |
| HVO_2284A | hypothetical protein | 2 | 0 |
| HVO_2254 | conserved hypothetical protein | 1 | 0 |
| HVO_2268 | TATA-binding transcription initiation factor | 1 | 0 |

But as stated below, none of the Halfvol4 proteins showed similarity to known viral proteins. However, the most abundant proteins identified were membrane-associated proteins such as ABC-transporters, S-layer protein, halocyanin, oxidoreductases, FtsZ and PilA. A similar observation was made upon analysis of extracellular vesicles of *Hfx. volcanii*; isolation of extracellular vesicles and of the pleolipovirus HFPV-1 in *Hfx. volcanii* and subsequent identification of proteins in these fractions revealed the presence of many membrane related proteins (Mills et al., 2024; Alarcon-Schumacher et al., 2022). Since our fractions were obtained from a strain with reduced EV production (Δ*oapA*) these cellular proteins very likey are derived from cell debris present in the culture supernatant.

## Discussion

Four putative provirus regions have been identified in the *Hfx. volcanii* genome (Halfvol1, Halfvol2, Halfvol3, and Halfvol4), which have not been extensively studied previously. Our research focused on characterising these proviruses, investigating their potential activity, and understanding their impact on *Hfx. volcanii*.

### The provirus deletion strain is phenotypically different from a provirus-containing strain

Since we could delete all four provirus regions, none of them has evolved to an essential part of the genome under the growth conditions used in the laboratory. A phenotypic comparison between the strain containing proviruses (H100) and the provirus-free strain (ΔHalfvol1-4) revealed various differences across several physiological parameters. ΔHalfvol1-4 showed a slightly faster growth compared to the control (Fig. 1 and Suppl. Fig. 3) under the majority of the conditions tested. We suggest that the growth enhancement observed in the provirus-free strain results from a reduction in the energy and resources required by *Hfx. volcanii* to replicate and transcribe the provirus genomes. Additionally, the production of viral particles could lead to the reduced growth of the provirus containing strain. A recent study identified a natural *Haloferax* strain (48N) closely related to the model organism *Hfx. volcanii* (Turgeman-Grott *et al*, 2024). This strain was chronically infected by a lemon-shaped virus. Deleting this virus from 48N resulted in significant alterations in its gene expression profile and a substantial increase of its growth rate (Turgeman-Grott *et al*., 2024).

In addition, ΔHalfvol1-4 exhibits higher resistance to oxidative stress in comparison to the control (Fig. 4), while the UV stress test showed no difference between the deletion strain and control strain. Hydrogen peroxide (H_2_O_2_) is a reactive oxygen species (ROS) that can cause significant damage to cells, including DNA, proteins, and lipids, leading to broad-ranging damage. In contrast, UV radiation primarily induces damage to DNA. Therefore, provirus genes seem to interfere with resistance against ROS. Virus infection is known to induce oxidative stress in the infected cells (Ivanov *et al*, 2017; Su *et al*, 2013). ROS response might include increased activity of virus defence systems, therefore it would be beneficial for a virus to block or reduce oxidative stress response. Which genes are responsible for this will be subject of future analyses.

Furthermore, ΔHalfvol1-4 cells display an elongated shape (Fig. 2), a characteristic often associated with motility in bacteria and archaea (Schiller *et al*, 2024; Schwarzer *et al*, 2021) suggesting that the proviruses have an influence on the host motility machinery. A correlation between the motility and oxidative stress response has been discovered during interaction between *Campylobacter jejuni* and its virus NCTC 12673 (Sacher *et al*, 2021).

### Proviruses affect motility and CRISPR-Cas expression providing thereby a potential superinfection exclusion mechanism

RNA-seq analysis showed upregulation of archaella, chemotaxis, and pili genes in ΔHalfvol1-4, which are all involved in motility (Table 3). This result provides a molecular explanation for the hypermotility of ΔHalfvol1-4 observed in the swarming assay (Fig. 3). Numerous instances in the literature highlight the influence of viruses or proviruses on cellular motility behaviour and the expression of archaella. For instance, transcriptome analysis of *Hfx. volcanii* infected with the pleolipovirus HFPV-1 revealed an overall down-regulation of archaella genes (Alarcón-Schumacher *et al*., 2022). Similarly, the deletion of the prophage CP4-57 from *E. coli* induced motility through flagella expression, as evidenced by transcriptome analysis and swarming assay results (Wang *et al*., 2009). Additionally, Brathwaite and colleagues reported impaired flagellar motility in *Campylobacter* due to the presence of the phage carrier state (Brathwaite *et al*., 2015).

Proviruses also seem to affect the immune system of *Hfx. volcanii*. RNA-seq analysis reveals a significant downregulation of CRISPR arrays (−3 or −2 log2FC) in the strain lacking proviruses, suggesting an interplay between proviruses and the CRISPR-Cas virus defence system (Table 3). Interestingly, Zegans et al. observed that a functional CRISPR-Cas system is necessary to inhibit swarming motility of *P. aeruginosa* by the DMS3 prophage (Zegans *et al*, 2009).

Proviruses frequently encode superinfection exclusion mechanisms. We propose that the proviruses present in *Hfx. volcanii* likely confer protection against other viruses through such mechanisms. Our hypothesis is based on the observation that the wild type strain containing proviruses exhibits higher expression of crRNAs compared to the provirus-free strain, as revealed by RNA sequencing data. This indicates that the presence of proviruses enhances the immune response by increasing crRNA production, thereby protecting *Hfx. volcanii* against other invading viruses. Furthermore, it appears that the presence of proviruses suppresses the expression of archaella/pili genes in *Hfx. volcanii*. This inhibition could prevent other viruses from entering the host cells, if they rely on these structures for attachment and adsorption. Numerous viruses utilise flagella or pili for cellular entry (Tittes *et al*, 2021b). For instance, bacteriophage x has been observed to attach to the flagella of *E. coli* via a tail fibre, subsequently migrating to the base of the flagella to inject its DNA (Schade *et al*, 1967). Similarly, ΦCB13 and ΦCbK viruses, infecting *Caulobacter crescentus*, adhere to the host’s flagella through a filament on the phage head (Guerrero-Ferreira *et al*, 2011), while phage 7-7-1, infecting *Agrobacterium sp*. strain H13-3, utilises tail fibres to attach to the flagella for absorption (Lotz *et al*, 1977).

In this context, proviruses within *Hfx. volcanii* may downregulate archaella expression as a mechanism to prevent other viruses from entering and infecting the host. This dynamic hints at a potential mutualistic relationship, wherein lysogeny benefits the chronic virus by preventing further infections from competing viruses, while simultaneously increasing the host protection against more virulent viruses through superinfection exclusion mechanisms. Moreover, the observed reduction in host motility, often perceived as negative, could serve as a strategy for self-quarantine: by restricting its own movement, *Hfx. volcanii* minimises virus spread and thereby reducing the chance of virus transmission. This may be a general strategy employed by hosts infected by a persistent virus (Chung *et al*, 2014).

Additionally, the Halfvol4 region contains several defence systems: a restriction modification system (rmeR, rmeM, rmeS) and the Avast2 protein (Suppl. Fig. 1), thus enhancing the protection of *Hfx. volcanii* against other invading viruses.

The superinfection exclusion effect of the provirus Halfvol3 against the pleolipovirus HFPV-1 has been documented (Alarcón-Schumacher *et al*., 2022). It was shown that the strain ΔHalfvol3 is more susceptible to HFPV-1 compared to the control. This suggests that the provirus region likely harbours one or more genes involved in mitigating the impact of HFPV-1. Moreover, in this study, the infection of HFPV-1 in a wild type *Hfx. volcanii* strain led to a notable downregulation of genes of Halfvol1. Specifically, VP3 was suppressed over 150-fold compared to uninfected cultures, while VP4, HVO_0268, and HVO_0270 exhibited decreases in expression with fold changes ranging between 40 and 60. These findings suggest that Halfvol1 may trigger a defensive response against HFPV-1, highlighting the intricate interplay of virus-virus interactions within the host (Alarcón-Schumacher *et al*., 2022).

Chronic infection is widespread within the archaeal domain, particularly among haloarchaea. It has been suggested that in archaea, chronic infection serves as a strategic trade-off in which the host cell maintains a persistent viral infection to protect itself from a more dangerous virus that could otherwise kill it (Turgeman-Grott *et al*., 2024; Wirth & Young, 2020).

Given the rarity of *Hfx. volcanii* viruses and the knowledge that proviruses often carry genes for superinfection exclusion mechanisms aimed at safeguarding the host from other viral threats, it is conceivable that the four proviruses present within *Hfx. volcanii* confer some level of superinfection exclusion. Consequently, the provirus-free strain may serve as a more susceptible host to viral infections, a factor with potential implications for future virologic research in *Hfx. volcanii*.

### Active viruses are generated by provirus circularisation and excision

Our results confirm previous data that Halfvol1 and Halfvol2 can be activated in *Hfx. volcanii* (Dyall-Smith *et al*., 2021). Excision of viral DNA and its presence in the supernatant were confirmed through PCR and Sanger sequencing. Specifically, circular DNA of Halfvol1, Halfvol2, and Halfvol3 (*attP* site) was detected in the culture supernatant (Fig. 5). Further evidence of provirus excision is provided by generation of PCR products across the chromosomal *attB* site, confirmed by sequencing (Fig. 7). This demonstrates that *Hfx. volcanii* strains possess chromosomal copies lacking proviruses and in addition confirms the spontaneous excision of proviruses from the chromosome, suggesting a persistent infection mode for these viruses.

Provirus activation was predominantly observed when cultures are grown at 30°C, suggesting that these viruses were more active at temperatures below the host’s optimal growth temperature (45°C) (Fig. 6). There are several documented instances where viruses exhibit increased activity at lower temperatures. For example, the archaeal pleomorphic virus AvPV1 shows higher titres at lower temperatures (Baquero *et al*, 2024), and the pleolipovirus HFPV1 was isolated from *Hfx. volcanii* culture grown at 28°C (Alarcón-Schumacher *et al*., 2022). It was demonstrated that elevated temperatures can impact phage infectivity by altering the structural integrity and flexibility of their lipid membranes or capsid proteins, which may influence their lifecycle strategies (Zhang *et al*, 2022). Temperature shifts are known to potentially trigger viral activity (Choi *et al*, 2010; Makky *et al*, 2021; Schuster *et al*, 1973; Zhang *et al*., 2022), indicating a possible ecological interaction between host and virus during colder periods. Numerous studies suggest that phage life cycle strategies may vary with the seasons. For instance, research in Antarctic Salt Lake and Tampa Bay has reported a higher prevalence of lysogeny during winter and spring compared to summer (Zhang *et al*., 2022). It is important to note that seasonal variations involve changes in several environmental factors, such as temperature, salinity, and primary productivity, highlighting the complex interplay of elements that influence phage lifecycle dynamics on a seasonal basis (Zhang *et al*., 2022).

Electron microscopy analysis of virus enriched fractions from the EV-reduced strain revealed that the first band contained, besides residual extracellular vesicles, structured particles which were quite large (app. 80 nm in diameter) and were not observed in ΔHalfvol1-4 (Fig. 9A). A collection of such particles illustrates that they are quite uniform and thus are candidates to represent pleolipovirus structures (Suppl. Fig. 7). The second band contained head-tailed structures which were not observed in ΔHalfvol1-4 (Fig. 9B, Suppl. Fig. 7). It is yet unresolved if the “tails” represent virus tails or if they represent cellular surface structures to which virus particles have attached. Thus, the particles isolated from the supernatants of Δ*oapA* cultures might be activated proviruses of *Hfx. volcanii.* This hypothesis is supported by the considerable differences observed between the Δ*oapA* and ΔHalfvol1-4 samples by TEM. While small and heterogenous particles are found in both samples, the larger and more uniform particles were observed exclusively in the Δ*oapA* strain. Additionally, mass spectrometry analysis of the CsCl density fractions of Δ*oapA* revealed the presence of viral proteins (Table 5).

The head-tailed structure observed in the Δ*oapA* supernatant (Figure 10B.) could be viral particles (pleolipovirus encoded by Halfvol1 or Halfvol 3, or the novel group virus encoded by Halfvol2) associating with fragments of archaella/pili or some other type of filament from *Hfx. volcanii*. According to this hypothesis, the virus would bind to archaella or pili during the adsorption step to facilitate entry into the host cell.

Taken together, the results indicate that *Hfx. volcanii* proviruses are active and replicate at low level, causing a type of persistent infection.

### Is Halfvol4 a provirus or an integrated plasmid?

Halfvol4 was initially described as a provirus (Hartman *et al*., 2010; Norais *et al*., 2007), however, upon closer examination of the Halfvol4 genome, it was not possible to detect any candidates for capsid proteins. Also, none of the proteins showed significant similarity to viral proteins. Halfvol4 exhibits characteristics of integrative and conjugative elements (ICEs). Interestingly, Halfvol4 encodes a T4SS, which functions similarly to plasmid conjugation systems, facilitating the transfer of ICEs between cells (Johnson & Grossman, 2015). ICEs range in size from 18 kb to 500 kb and are typically AT-rich compared to their host genome. They often carry cargo genes that provide phenotypic traits to host cells, such as antibiotic resistance, pathogenesis, restriction modification (as also seen in Halfvol4), and carbon-source utilisation (Johnson & Grossman, 2015). It is important to note that ICEs are usually flanked by *att* sites and form circular DNA upon excision. A slightly extended Halfvol4 (as compared to its original description) is enclosed in a 14 nt direct repeat which resembles an *att* site. An additional copy of the 14 nt sequence is encountered internally within Halfvol4. Halfvol4 appears to be inserted in the *Hfx. volcanii* genome sequence when compared to some closely related haloarchaeal strains that do not have an insertion at this position. A few other strains have inserts at the corresponding position however these inserts have a different sequence. In *Hfx. alexandrinus* pws11, the insert shows similarity to Halfvol4, in *Haloferax* sp. Atlit-105R the insert is totally unrelated (Suppl. Table 2). Circular Halfvol4 DNA was not detected in the cell fraction or culture supernatant. Therefore, we suggest that Halfvol4 is not a provirus but a plasmid that has been integrated into the *Hfx. volcanii* genome.

## Methods

### Strains and growth conditions

Strains are listed in Suppl.Table 3. *Hfx. volcanii* strain H100 was used as control strain for all experiments. In short, it is derived from wild-type strain DS2 by curing of plasmid pHV2 and deletion of three genes that can be used as selection markers (Δ*pyrE2*, Δ*leuB,* Δ*hdrB*) (Allers *et al*, 2004). Strain H100 was grown aerobically with shaking (200 rpm) at 45°C in Hv-YPCab medium supplemented with thymidine. The parental strain of ΔHalfvol1-4 is H133 (Δ*pyrE2* Δ*leuB* Δ*hdrB* Δ*trpA*).

We use strain H100 (Δ*pyrE2* Δ*leuB* Δ*hdrB*) as a control which differs from strain H133 by carrying the *trpA* gene. This is consistent with the *trpA* gene being present in ΔHalfvol1-4 at the position of Halfvol3. However, while the *trpA* gene is under control of its native promoter in strain H100, it is under the control of p.fdx in ΔHalfvol1-4. Strain Δ*oapA* was previously described (Mills *et al*., 2024; Wolters *et al*, 2019), its parent strain is H26 (Δ*pyrE2*) (Allers *et al*., 2004). Strain ΔHalfvol1-4 was generated in the current study (see below).

### Cloning of plasmids

Plasmids and primers are listed in Suppl.Table 3. Plasmid pTA1102 (also termed pTA131-UP-DO (Halfvol4)) is a pTA131 plasmid with insertion of 1,504 bp *Hfx. volcanii* genomic DNA fragment containing downstream flanking region of Halfvol4 (bp 2163565-2165068), and 3,100 bp *Hfx. volcanii* genomic DNA fragment containing upstream flanking region of Halfvol4 (bp 2108260-2111359). The region deleted is bp 2111359-2163565, or HVO_2252-2293. To generate pTA131-UP-DO (Halfvol1), first the upstream region of Halfvol1 was amplified by PCR using primers Provir6-up.fw and Provir6-up.rev. Plasmid pTA131 as well as the insert were digested with *Eco*RV and ligated together to generate pTA131-UP (Halfvol1). The downstream region of Halfvol1 was also amplified by PCR using primers ProVir6-do.fw and ProVir6-do.rev. The resulting fragment as well as plasmid pTA131-UP (Halfvol1) were digested with *Eco*RV and ligated to generate plasmid pTA131-UP-DO (Halfvol1). To generate pTA131-UP-DO (Halfvol2), the upstream region of Halfvol2 was amplified by PCR using primers Up KpnI Halvol2 F and Up HindIII Halvol2 R. Plasmid pTA131 and insert were digested with *Kpn*I and *Hind*III and ligated together to generate pTA131-UP (Halfvol2). The downstream region of Halfvol2 was generated by PCR using primers Do HindIII Halvol2 F and Do XbaI Halvol2 R. The resulting fragment as well as plasmid pTA131-UP (Halfvol2) were digested with *Hind*III and *Xba*I and ligated together to generate plasmid pTA131-UP-DO (Halfvol2). pTA231-p.fdx-Nflag-bridgCDS Halfvol1 plasmid was obtained by amplifying the insert using primers CDS1-SnaBI F and CDS1-XbaI R. The resulting fragment and the pTA231-p.fdx-Nflag plasmid were subjected to digestion with *SnaB*I and *Xba*I, followed by ligation. pTA231-p.fdx-Nflag-bridgCDS Halfvol2 plasmid was obtained by amplifying the insert using primers CDS2-SnaBI F and CDS2-XbaI R. The resulting fragment and the pTA231-p.fdx-Nflag plasmid were subjected to digestion with *SnaB*I and *Xba*I, followed by ligation. pTA231-p.fdx-Nflag-bridgCDS Halfvol3 plasmid was obtained by amplifying the insert using primers CDS3-SnaBI F and CDS3-XbaI R. The resulting fragment and the pTA231-p.fdx-Nflag plasmid were subjected to digestion with *SnaB*I and *Xba*I, followed by ligation.

### Deletion of proviruses

The provirus-free strain was generated using the pop-in pop-out method as previously described (Bitan-Banin *et al*, 2003). A Halfvol3 deletion strain was obtained from Uri Gophna (Tel Aviv University). In this strain, the Halfvol3 provirus is replaced by the *trpA* gene, which is controlled by the p.*fdx* promoter. The presence of the *trpA* marker at the position of Halfvol3 was confirmed by sequencing (Alarcon-Schumacher, personal communication), but was inadvertently not reported in the original publication (Alarcón-Schumacher *et al*., 2022). The Halfvol4 locus was deleted in the ΔHalfvol3 strain using the deletion plasmid pTA1102 (pTA131-UP-DO (Halfvol4)). Subsequently, Halfvol1 was deleted in the ΔHalfvol3ΔHalfvol4 strain using the plasmid pTA131-UP-DO (Halfvol1) and lastly, the provirus Halfvol2 was removed from the ΔHalfvol1ΔHalfvol3ΔHalfvol4 strain using the plasmid pTA131-UP-DO (Halfvol2). This resulted in strain ΔHalfvol1-4 which is devoid of all proviruses.

### Determination of growth curves under normal and stress conditions

Growth curves were generated using 96-well microtiter plates and the Epoch 2 plate reader (BioTek Instruments). Cultures of strains H100 and ΔHalfvol1-4 were diluted to an OD_650_ of 0.05, and 200 μl of this diluted culture was added to the 96-well plates in triplicate or quintuplicate, along with a control containing only the Hv-YPCab supplemented thymidine medium. Media with 18% BSW were used for the standard salt condition, while 15% BSW and 23% BSW were used for low and high salt stress conditions, respectively. Three biological replicates were performed. The standard temperature was 45°C, with 30°C and 50°C applied for temperature stress conditions. To minimise evaporation, 300 μl of medium was added to the outer wells of the plate. The optical density was recorded at 650 nm at 30-minute or 1-hour intervals (Dyall-Smith, 2009). The growth data were analysed using Excel.

### H_2_O_2_ survival assay

*Hfx. volcanii* was inoculated into 4ml Hv-YPCab medium with thymidine and incubated at 45°C with shaking at 200 rpm. When the cultures reached an OD_650_ of 0.3 to 0.4, four 480 µL aliquots were taken and transferred into 2 ml reaction tubes. To these samples, H_2_O_2_ stock solution was added to achieve final concentrations of 2 mM, 4 mM, and 6 mM H_2_O_2_, resulting in 500 µl cell suspensions. For the control, 20 µl of 18% SW was added instead of H_2_O_2_. Samples were incubated for 1h at 45°C with shaking at 450 rpm. Serial dilutions were prepared using 18% SW, ranging from 10^−3^ to 10^−6^, and 20 µl of each dilution was spotted in triplicate onto pre-warmed Hv-YPC + thymidine agar plates and droplets were allowed to absorb. Plates were incubated for 3-4 days at 45°C. Colonies were counted, and survival fractions were calculated by averaging the triplicates for the 10^−5^ dilution level.

### UV survival assay

Cultures were grown to an OD_650_ of approximately 0.4. Cells were then serially diluted (10^0^-10^−8^) in 18% BSW. Quadruplicate 20 μl samples were spotted onto Hv-YPC + thymidine plates, one plate for each exposure time (including a non-illuminated control). The plates were air dried for 30 min to allow the spotted culture to dry out. The plates were irradiated for 20, 40, 60, 80 and 100 sec with a wave length of 254 nm. After irradiation, plates were incubated in a black bag at 45°C for 4 days. Following the incubation period, the colonies were counted.

### Light microscopy

Cultures with OD_650_ values of 0.3 (early exponential phase), 0.6 (late exponential phase), and 1.0 (stationary phase) were used. Cells were concentrated by centrifuging 1 mL of preculture at 4,000 x g for 6 minutes at room temperature. The supernatant was discarded, and the cell pellet was resuspended in 100 to 200 μl Hv-YPCab + thymidine medium. For microscopy, slides were prepared with a thin agar layer by boiling 0.1 g of agarose with 4 mL of ddH_2_O and 6 mL of 30% saltwater, then mixing the solutions. A thin agarose bed was applied to a slide with a brush, briefly dried, and 3-5 μl of cell suspension was added. The sample was covered with a coverslip and sealed with clear nail polish. Microscopy was performed using a Leica DM5500 B light microscope in phase contrast at 100x magnification with oil immersion.

### Motility analysis

To create soft agar plates (0.33%), 1 g of micro-agar was dissolved in 100 mL of demineralized water. After fully dissolving the agar in a microwave oven, 200 ml of pre-warmed 30% BSW was added with gentle swirling. The mixture was autoclaved for sterilization and then placed on a heated magnetic stirrer plate, where 34 ml of 10x Hv-Ca and and other components (CaCl_2_, thymidine and thiamine & biotin) were added as required. The liquid medium was poured into large plates, allowed to solidify for 1-2 hours, and cultures were inoculated deeply into the semi-liquid medium with a toothpick. Three replicates for each strain were spotted in the same plate. Plates were incubated upside down at 45°C for 4-5 days to observe swarming behaviour.

### Biofilm formation

Cultures were grown until late exponential phase, and 150 µl were added to each well of a microtiter plate, with 5-10 replicates and a negative control. Plates were incubated without shaking at 45°C and 30°C for 72 hours. After incubation, liquid culture was removed, wells were washed twice with 18% BSW, fixed with 2% acetic acid, and stained with 0.1% crystal violet. Plates were washed three times with ddH_2_O and air-dried. Finally, wells were destained with a solution of 10% acetic acid and 30% ethanol, and OD_600_ was measured in a microtiter plate photometer.

### RNA-seq analysis

Three biological replicates of the control strain (H100) and the deletion strain (Δ1-4) were grown at 45°C and 200 rpm until 0.5 OD_650_ was reached. Then, 20 ml of culture was centrifuged at 4°C and 6,000 x g for 10 minutes. The supernatant was discarded and the cell pellet was stored at −80°C until further use. For RNA preparation, the thawed pellet was resuspended in 1 ml NucleoZOL™ (Macherey-Nagel) and after the addition of 400 µl ddH_2_O, the mixture was vortexed for 30 sec and incubated at room temperature for 10 min to lyse the cells. After centrifugation at 13,000 x g for 15 min at room temperature, the upper phase was carefully transferred to a new tube and 5 µl of 4-bromanisol were added, followed by vortexing and incubation for 10 min. The tubes were centrifuged at 13,000 x g for 15 min at room temperature. The supernatant was transferred to a new tube and mixed with 1 ml of isopropanol. After incubation for 10 min at room temperature, the RNA was pelleted by centrifugation at 14,000 rpm for 10 min at 4°C. The pellet was washed twice with 70% ethanol and then resuspended in 240 µl of RNase-free ddH_2_O.

To remove DNA, 30 µl of RQ1 DNase buffer and 30 µl of RNase-free RQ1 DNase were added and the mixture was incubated at 37°C for 1 h. RNA was then precipitated by ethanol and resuspended in 100 µl ddH_2_O. Since *Hfx. volcanii* is a polyploid organism with a large amount of DNA, the samples were subjected to another step of DNase digestion using the TURBO™ DNase digestion according to the manufacturer’s instructions. After 30 min incubation at 37°C, phenol-chloroform extraction was performed followed by ethanol precipitation. The RNA pellet was dissolved in 50 µl of RNase-free ddH_2_O and the concentration determined using a NanoPhotometer^®^ N60 (Implen). 10 µg of RNA samples were further processed at the Core Unit Systems Medicine facility at the University of Würzburg. Samples underwent depletion of ribosomal RNA molecules using a commercial Pan-Archaea riboPOOL rRNA depletion kit (siTOOLs Biotech, dp-K024-000027). The ribo-depleted RNA samples were first fragmented using ultrasound (4 pulses of 30 s at 4°C). Then, an oligonucleotide adapter was ligated to the 3’ end of the RNA molecules. First-strand cDNA synthesis was performed using M-MLV reverse transcriptase with the 3’ adapter as primer. After purification, the 5’ Illumina TruSeq sequencing adapter was ligated to the 3’ end of the antisense cDNA. The resulting cDNA was PCR-amplified using a high-fidelity DNA polymerase and the barcoded TruSeq-libraries were pooled in approximately equimolar amounts. Sequencing of pooled libraries, spiked with PhiX control library, was performed at 13-16 million reads per sample in single-end mode with 100 cycles on the NextSeq 2000 platform (Illumina). Demultiplexed FASTQ files were generated with bcl-convert v4.0.3 (Illumina). Raw sequencing reads were subjected to quality and adapter trimming via Cutadapt (Martin, 2011) v2.5 using a cutoff Phred score of 20 and discarding reads without any remaining bases (parameters: --nextseq-trim=20 -m 1 -a AGATCGGAAGAGCACACGTCTGAACTCCAGTCAC). Afterwards, all reads longer than 11 nt were aligned to the Haloferax volcanii DS2 reference genome (RefSeq assembly accession: GCF_000025685.1 excluding plasmid pHV2 (NC_013965.1)) using the pipeline READemption (Förstner *et al*, 2014) 2.0.3 with segemehl v0.3.4 (Hoffmann *et al*., 2009) and an accuracy cut-off of 95% (parameters: -l 12 -a 95). READemption gene_quanti was applied to quantify aligned reads overlapping genomic features by at least 10 nt (-o 10) on the sense strand based on annotations (CDS, mature_transcript, rRNA, tRNA) originating from a detailed continuous manual curation effort (Laass et al, 2019). The annotation version from 28-MAR-2023 was used (annotated protein sequences in: https://doi.org/10.5281/zenodo.7794769), this version includes small ORFs reported in (Hadjeras et al., 2023). Based on these counts, differential expression analysis was conducted via DESeq2 (Love *et al*., 2014) v1.24.0. Read counts were normalized by DESeq2 and fold-change shrinkage was conducted by setting the parameter betaPrior to TRUE. In addition, outlier flagging by Cook’s distance was disabled in the results function (cooksCutoff=FALSE). Differential expression was assumed at adjusted p-value after Benjamini-Hochberg correction (padj) < 0.05 and |log2FoldChange| ≥ 1.

### Methods used for virus purification and concentration

1 l cultures of strains Δ*oapA* and ΔHalfvol1-4 with an OD_650_ of 0.05 were incubated at 30°C with shaking at 180 rpm until reaching the stationary phase (OD_650_ 0.86 – 1.0). The culture was centrifuged at 5,524 x g at 4°C for 30 min. The supernatant, potentially containing viral particles, was subjected to another centrifugation step to eliminate any remaining cells. To precipitate the virus, 1:4 volume of cold 40% PEG6000 was added to the supernatant (final concentration 10%). After overnight incubation at 4°C, the viral suspension was centrifuged at 13,500 x g for 1 h at 4°C, and the pellet was resuspended in 10 ml of 18% BSW. Further purification involved three filtration steps: the first with a 0.45 µm pore-size filter, followed by two with 0.22 µm filters. Another precipitation step with cold 40% PEG6000 was performed, followed by overnight incubation at 4°C. After centrifugation at 15,000 x g for 1 h at 4°C, the supernatant was removed, and the pellet was resuspended in 1 ml of 18% BSW. To eliminate free nucleic acids, 10 µl of RNase A and 20 µl of DNase (1000 U) were added to the sample, which was then incubated for 2 h at 37°C. The digested samples were either stored at 4°C or immediately processed for CsCl density gradient centrifugation.

A CsCl solution, having a density of 1.2 g/ml and a concentration of 0.342 g/ml in 18% BSW, was filtered through a 0.22 µm filter. Subsequently, 11 ml of this solution was dispensed into Ultra-Clear™ centrifuge tubes (14 x 95 mm), followed by gently layering 1 ml of the viral sample over the CsCl solution. The tube was centrifuged at 38,000 x g for 21-24 h at 4°C using the SW40T1 rotor in the Optima L-60+ Ultracentrifuge (Beckman Coulter). After ultracentrifugation, the bands were collected by carefully aspiring 1 ml fractions of the gradient from the top. Each fraction was supplemented with 10% PEG6000 (final concentration) and incubated overnight at 4°C. Following centrifugation at 15,000 rpm for 1 h at 4°C, the pellet was resuspended in 50 to 100 µl of 18% BSW. To analyse for the presence of viral genomes, PCR analysis was conducted using 1 µl of a 1:10 diluted sample as the template. A total of 30 or 31 cycles was applied.

### Detection of provirus excision by PCR analysis

To detect virus DNA circularisation, a PCR was performed using divergent primers located at the ends of the provirus region (illustrated in Fig. 5). The primers employed for the detection of circular Halfvol1 are prov6 circ R and prov6 circ F; for Halfvol2, Prov2 circ F and Prov2 circ R were used; and for Halfvol3, prov5 circ F and prov5 circ R were employed (Suppl. Table 3C.). The PCR mixture included 5 µL of 5x Green GoTaq Reaction Buffer, 1.25 µL of DMSO, 0.5 µL of dNTPs (5 mM), 0.25 µL each primer (500 ng/µL), 1 µL of template (a 1:10 dilution of isolated virus particles from the culture supernatant to avoid salt interference), 17.5 µL of ddH_2_O and 0.25 µL of Go-Taq polymerase, for a total reaction volume of 25 µL. The initial denaturation was carried out at 95°C for 5 minutes, followed by 31 cycles of denaturation at 95°C for 30 seconds, annealing temperature at 62°C for 30 seconds, and extension at 72°C for 30 sec. After cycling, a final extension was performed at 72°C for 3 minutes. PCR products were analysed on a 1% agarose gel. The resulting bands were excised and sent for Sanger sequencing.

### Transmission electron microscopy (TEM)

Samples from culture supernatants were fixed with a 2.5% glutaraldehyde solution in 18% BSW for 25 min. Approximately 7 µl of the fixed samples were placed onto glow-discharged 300 mesh copper grids covered with a carbon enhanced formvar film. After 10 min, excess liquid was removed by tapping the grid on filter paper soaked with ddH_2_O. The grids were then washed three times in droplets of ddH_2_O, with excess water removed after each wash. The sample was stained with a 2% aqueous uranyl acetate solution for 4 sec. After drying, the samples were imaged using a TEM (JEM1400, Jeol, Japan).

### Mass spectrometry

The second band derived from strain Δ*oapA* and the corresponding pink band from ΔHalfvol1-4 after CsCl density gradient centrifugation were concentrated by PEG precipitation (see above). The pellet was resuspended in approximately 100 µl of 50 mM Tris HCl pH 7.2. The samples were centrifuged at 10,000 x g for 10 min to remove insoluble particles, and protein concentration was measured using a Bradford assay (Roti-Nanoquant, Carl Roth GmbH). 10 µg of protein from each sample was loaded onto a 12% SDS gel and stained with Coomassie Brilliant Blue. SDS-PAGE-separated protein samples were processed as described by Shevchenko *et al*. (Shevchenko *et al*, 1996a; Shevchenko *et al*, 1996b). The resulting peptides were loaded to nano HPLC coupled to an Exploris Orbitrap Mass spectrometer (Thermo Fisher). The peptides were separated with a linear gradient of 5–40% buffer B (80% acetonitrile and 0.1% formic acid) at flow rate of 300 nL/min over 52 min total gradient time. The MS data was acquired by scanning the precursors in mass range from 350 to 1400 m/z at a resolution of 60,000 at m/z 200. Top30 precursor ions were chosen for MS2 by using data-dependent acquisition (DDA) mode at a resolution of 15,000 at m/z 200. Data analysis and search was performed using Maxquant Software (1.6.17.0) and Scaffold 5.2.1 with Uniprot_Haloferax_volcanii database from August 2021 with 4,186 entries.

### Pulsed Field Gel Electrophoresis

For PFGE, genomic DNA was prepared in agarose plugs as described previously (Delmas *et al*, 2009). For analysis of intact genomic DNA, agarose plugs were subjected to 100 Gy of γ radiation using a ^137^Cs source (Gammacell 1000), to linearize circular chromosomes (Beverley, 1989). PFGE was performed using a CHEF Mapper apparatus (Bio-Rad). DNA fragments were separated on a 1.2% agarose gel in 0.5× TBE (Tris, borate, EDTA) at 14 °C, with a gradient voltage of 6 V/cm, linear ramping, an included angle of 120°, initial and final switch times of 0.64 s and 1 min 13.22 s, respectively, and a run time of 20 h 46 min, as described previously (Ausiannikava *et al*, 2018). The gel was stained with ethidium bromide.

### qPCR

For sample preparation, 1 ml of culture was grown to OD_650_ 0.8 and centrifuged at 1,503 x g for 8 min at room temperature. The supernatant was filtered through a 0.22 µm filter to remove cells. Subsequently, 1 ml lysis buffer was added and ethanol precipitation was performed. After centrifugation, the pellet was resuspended in 300 µl of ddH_2_O. Real-time quantitative PCR (RT-qPCR) was performed using a qtower3 touch qPCR machine (Analytik Jena) in 96-well plate. The primers used for each target were qPCR Halfvol1 F2 and qPCR Halfvol1 R2 for Halfvol1; qPCR Halfvol2 F1 and qPCR Halfvol2 R1 for Halfvol2; qPCR Halfvol3 F2 and qPCR Halfvol3 R2 for Halfvol3 (Suppl. Table 3C). The qPCR mixture included 10 µl of qPCRBIO SyGreen Blue Mix Separate-ROX (PCR Biosystems), 0.8 µl each primer (10 µM), 6.4 µl of ddH_2_O and 2 µl of template (1:2 or 1:10 diluted sample), for a total reaction volume of 20 µl. The initial denaturation was carried out at 95°C for 2 minutes, followed by 40 cycles of denaturation at 95°C for 5 seconds, annealing/extension temperature at 60°C for 30 seconds. First primer efficiencies were assessed (Halfvol1: 0.88 Halfvol2: 0.98 Halfvol3: 0.87). The level of circularised viral genome was quantified using the absolute quantitative method (Harshitha & Arunraj, 2021). Standard curves were generated with plasmid pTA231-p.fdx-Nflag-bridgCDS Halfvol1, pTA231-p.fdx-Nflag-bridgCDS Halfvol2 and pTA231-p.fdx-Nflag-bridgCDS Halfvol3 to calculate the absolute concentration of circularised Halfvol1, Halfvol2 and Halfvol3, respectively (Suppl. Table 3B.). Each sample was analysed in triplicate for both technical and biological replicates.

## Data availability

Data from RNA-seq and mass spectrometry will be deposited in public databases and full supplementary data will be made available upon publication of the manuscript.

## Acknowledgments

We thank Manuela Weishaupt, Susanne Schmidt, Elena Katzowitsch and Panagiota Arampatzi for expert technical assistance as well as Jessika Jakubowski and Sandra Schreiber for help with generating the provirus deletion strain ΔHalfvol3ΔHalfvol4. In addition, we thank Lisa-Katharina Maier and Uri Gophna for constructive and fruitful discussions as well as for careful reading of the manuscript.

