## Supplementary material for "Provirus deletion from *Haloferax volcanii* affects motility, stress resistance and CRISPR RNA expression": Suppl. Table 1

<u>Supplementary Figures</u>

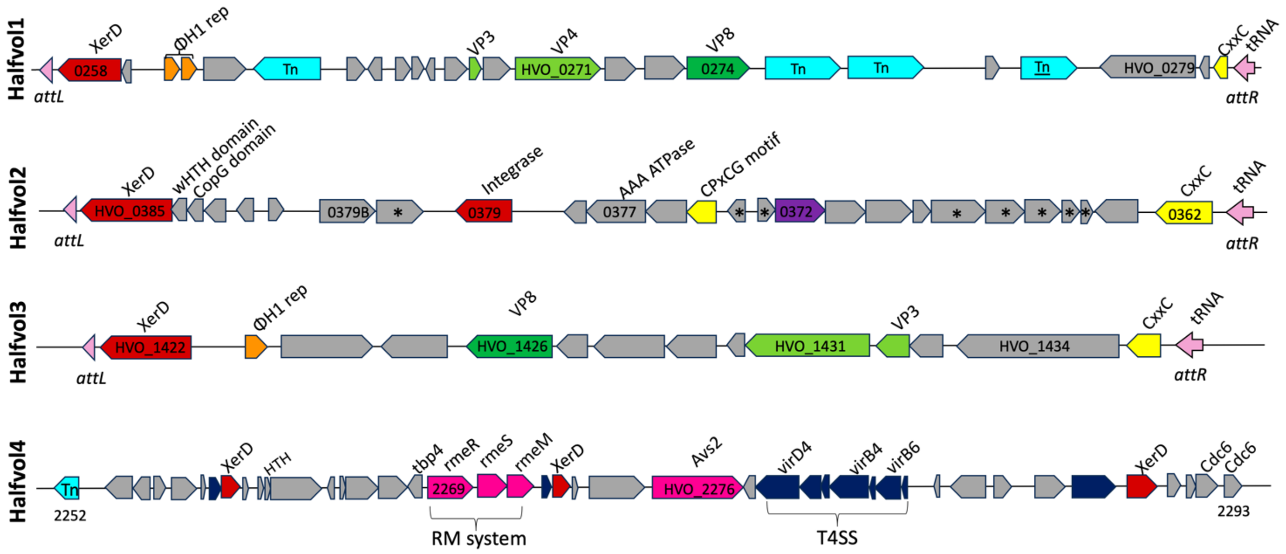

**Supplementary Figure 1. Genomic organisation of Halfvol1, Halfvol2, Halfvol3 and Halfvol4.**

The attachment sites (*attR* and *attL*) are marked by light pink triangles (partial tRNA) or arrow (complete tRNA gene) at the ends of Halfvol1, Halfvol2, and Halfvol3. Key features include the structural proteins VP3 and VP4 with light green arrows, the viral protein VP8 with a dark green arrow, the integrases (XerD and HVO_0379) in red, the CxxC motif protein/Zn finger protein in yellow and the HTH domain proteins, which are distantly related to homologues of the phiH1 repressor in orange. Light blue arrows indicate transposases (Tn), one of which is underlined to indicate the recombination event between chromosomal and pHV4 transposases, which allowed the integration of pHV4 into the main chromosome. The gene HVO_0372, which encodes the possible capsid protein for Halfvol2, is shown in purple (Dyall-Smith *et al*, 2021). Proteins with transmembrane domains in Halfvol2 are asterisked. In addition, fuchsia arrows highlight defence mechanisms: Avast type 2 (Avs2) and the restriction/modification (RM) system found in Halfvol4. The genes of Halfvol4 homologous to T4SS are shown in dark blue.

In Halfvol1, Halfvol2, and Halfvol3 the first proviral gene encodes an integrase while the last gene encodes a protein containing a CxxC motif (Dyall-Smith *et al.*, 2021). Recombination at the 14 bp direct repeat results in the circularisation and excision of the provirus, leaving the complete tRNA on the chromosome which serves as the *attB* site. The circularised viral DNA contains the 14 bp tRNA-derived region, which represents the *attP* site. The *attP* element plays a critical role in enabling the virus genome to integrate in a site-specific manner into the host chromosome through recombination with *attB*.

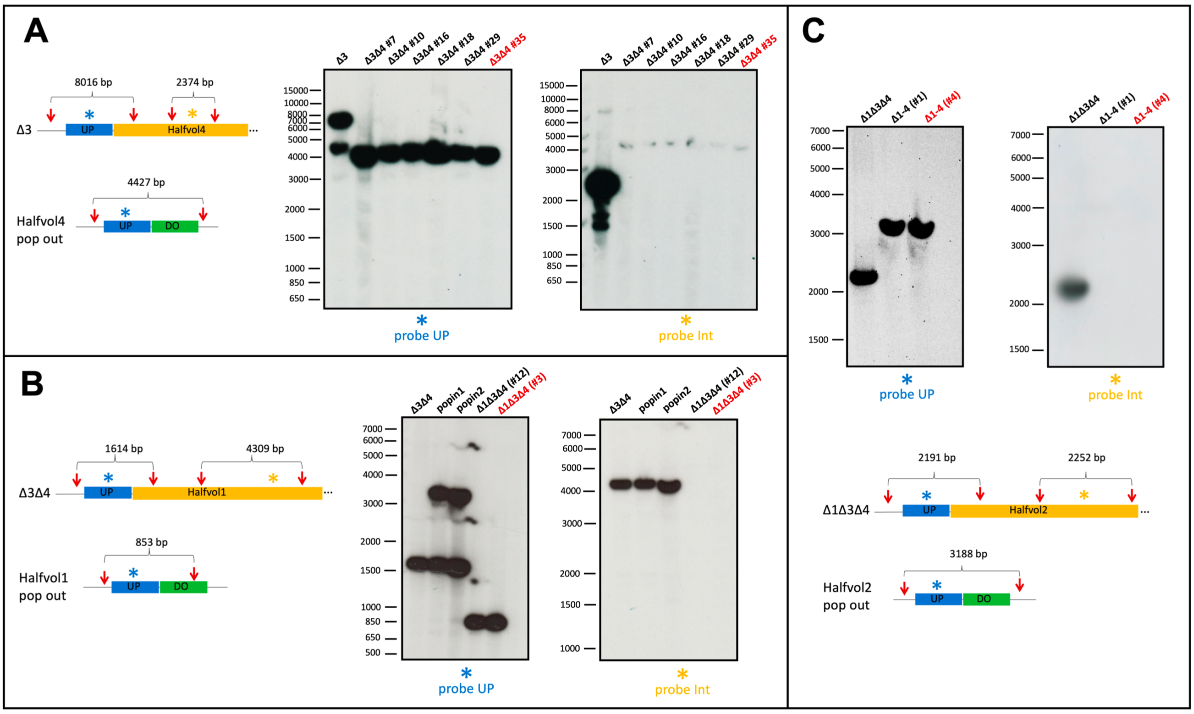

**Supplementary Figure 2. Southern blot analysis of provirus deletion strains.**

In each panel, a schematic drawing of the genomic location of wild type and deletion mutant are shown. Provirus regions are shown in yellow, labelled with the provirus name (not drawn to scale). Upstream (UP) regions are blue and downstream (DO) regions are green. For reasons of space, the downstream region is omitted in the wild type situation, when the complete provirus is still present. The approximate position of *Sal*I restriction sites (red arrows) and the size of relevant *Sal*I restriction fragments are indicated. The PCR probes used for hybridisation (300-500nt) are indicated by asterisks (upstream probe in blue; internal probe in yellow). Southern blots show gDNA samples after digestion with *Sal*I. Southern blots were hybridised with radioactively labeled PCR probes against the upstream region (probe UP) or against the provirus region (probe Int). Pop out clones, that have the provirus genome deleted, show a shorter *Sal*I fragment upon hybridisation with the upstream probe and no hybridisation with the internal probe. In each case, several pop out clones are shown. The pop out clone highlighted in red was selected for subsequent experiments. **A.** Deletion of Halfvol4 from the ΔHalfvol3 strain. **B.** Deletion of Halfvol1 from the ΔHalfvol3ΔHalfvol4 strain. **C.** Deletion of Halfvol2 from the ΔHalfvol1ΔHalfvol3ΔHalfvol4 strain.

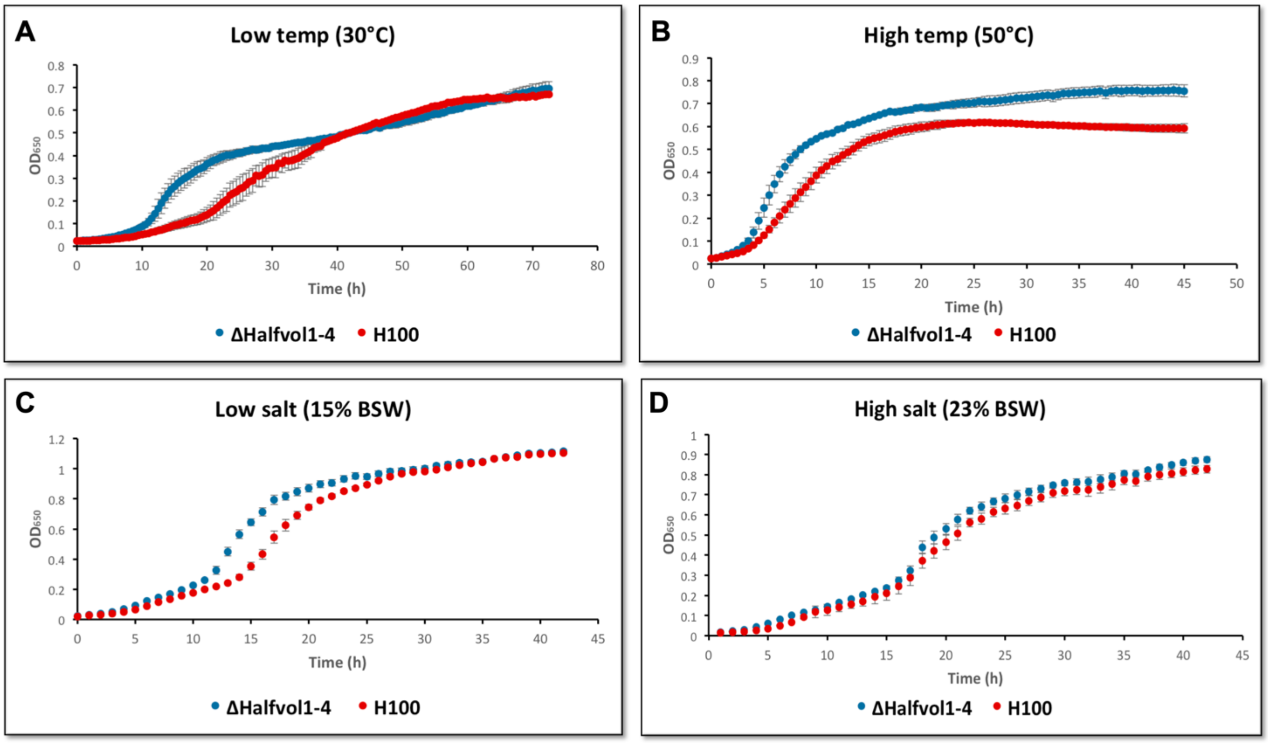

**Supplementary Figure 3. Growth curves of the provirus-free strain ∆Halfvol1-4 and the control strain H100 under stress conditions.**

Growth analysis of the provirus-free strain ∆Halfvol1-4 (in blue) and the control strain H100 containing the proviruses (in red) were performed at **A.** low (30°C) and **B.** high (50°C) temperatures, and at **C.** low (15% BSW) and **D.** high (23% BSW) salt concentration. See Fig.1 in the manuscript for growth under optimal conditions (45°C and 18% BSW).

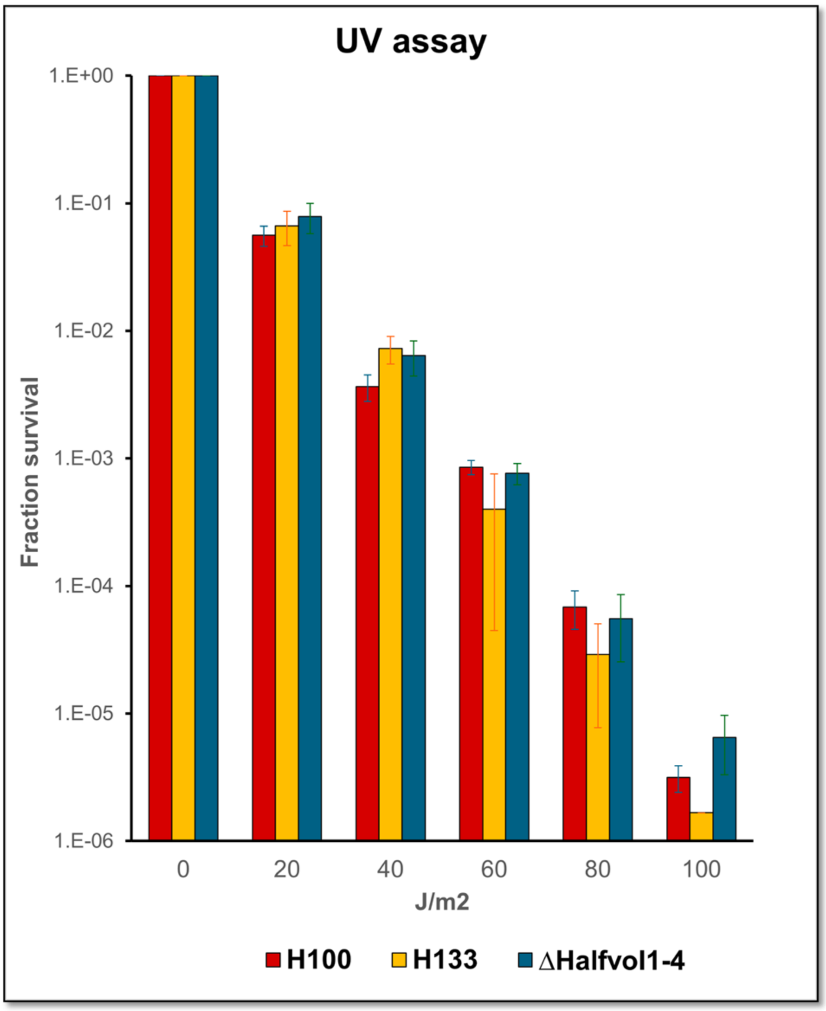

**Supplementary Figure 4. UV assay of the strains ΔHalfvol1-4, H100 and H133.**

The viability of the provirus-free strain ΔHalfvol1-4 (in blue), the control strain H100 (in red) and the parental strain H133 (in yellow) were tested after illumination with UV for different times. No significant differences were observed. The x-axis shows the amount of J/m2 used and the y-axis shows the fraction of survival in logarithmic scale.

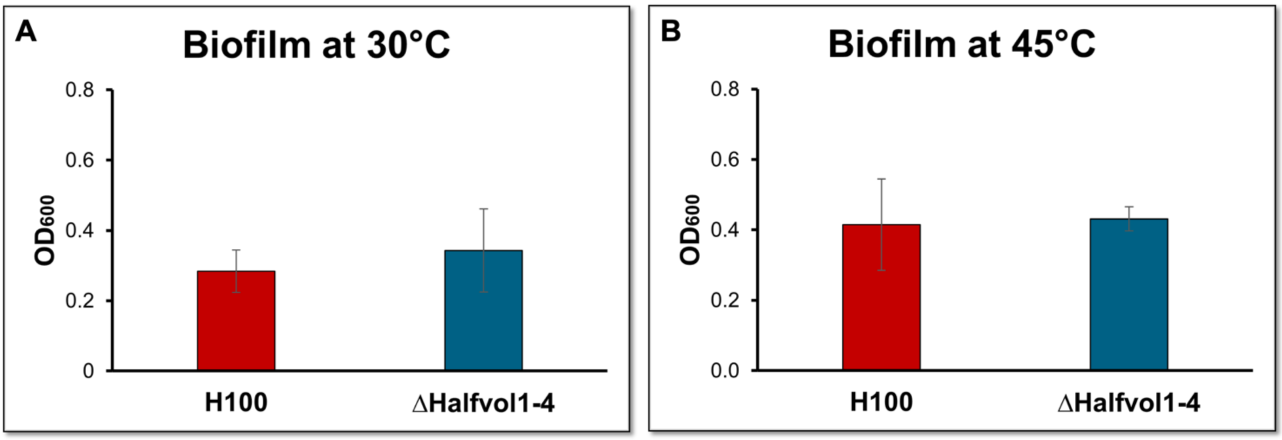

**Supplementary Figure 5. Biofilm assay of the provirus-free strain ΔHalfvol1-4 and the control strain H100 at 30°C and 45°C.**

Biofilm formation of the provirus-free strain ΔHalfvol1-4 (in blue) and the control strain H100 containing the proviruses (in red) were tested at 30°C (**A.**) and 45°C (**B.**). Absorbance was measured at 600 nm.

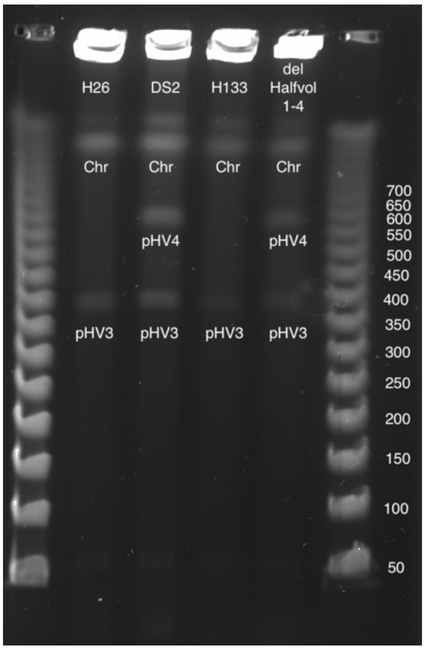

**Supplementary Figure 6. Pulsed field gel analysis confirms that pHV4 is present as an episome in the ∆Halfvol1-4 strain.**

Pulsed field gel electrophoresis was used to demonstrate that pHV4 has excised as an episome in the ∆Halfvol1-4 strain, whereas it is integrated on the chromosome in the parental strain H133 and in the laboratory strain H26; in the wild isolate DS2, pHV4 is originally present as an episome (Hawkins *et al*, 2013).

A. 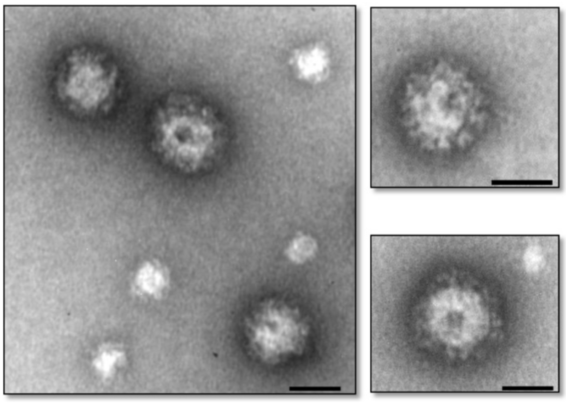

B. 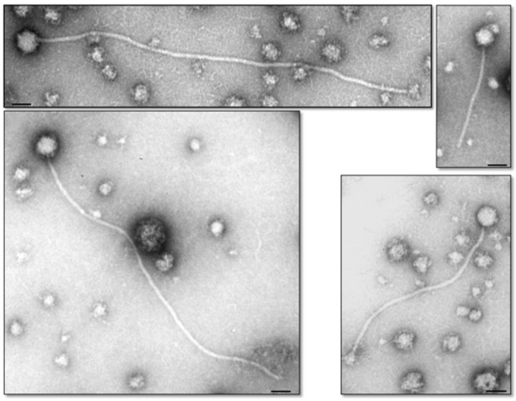

**Supplementary Figure 7. Collection of pleolipovirus-like and head-tailed structures found in the supernatant of ∆*oapA*.**

**A.** Collection of pleolipovirus-like structures, with the majority obtained from the first band. **B.** Collection of head-tailed structures, with the majority obtained from the second band. The scale bar of 50 nm is shown.

**Supplementary Table 1. Proviruses of *Hfx. volcanii*.**

This table lists the key characteristics of the four proviruses which have been reported for *Hfx. volcanii*. The extent of the virus is provided by position, the set of encoded ORFs, and size. All viruses are encode on the main chromosome. The direct repeat enclosing the provirus, which four all four proviruses is 14 nt long, is provided. For Halfvol1-3, only the *attP* is contained, thus excluding the repeat copy from *attB*. Thus, the provided length and region represents the circularised (activated) form of the provirus. For Halfvol2, all 14 nt of *attB* belong to the adjacent tRNA. For Halfvol1 and Halfvol3, 13 of the 14 nt of *attB* belong to the adjacent tRNA (letters in uppercase). Halfvol4 is not adjacent to a tRNA gene (repeat letters in lowercase). The boundaries of the provirus as reported in the literature may differ from those provided here. The morphotype "novel group" is from (Dyall-Smith *et al.*, 2021).

| **Name** | **Position (excluding *attB*)** | **ORFs (HVO_)** | **Size (nt)** | **Direct Repeat** | **Repeat Sequence** | **Morphotype** | **flanks (*attP*)** | **# inte-grases** | **Reference** |
|---|---|---|---|---|---|---|---|---|---|
| Halfvol1 | 231,452-252,024 | 0258 - 0280A | 20,573 | 14 nt | CCGGGCGGTCCCAt | betapleolipovirus | $tRNA^{Pro}$-TGG | 1 | (Bath *et al*, 2006; Dyall-Smith *et al.*, 2021) |
| Halfvol2 | 329,579-341,853 | 0362-0385 | 12,275 | 14 nt | TCCCCGCGAGTCCA | novel group | $tRNA^{Ala}$-CGC | 2 | (Dyall-Smith *et al.*, 2021) |
| Halfvol3 | 1,294,959-1,307,485 | 1422-1434A | 12,527 | 14 nt | CCTGTAGCGCCCAt | alphapleolipovirus | $tRNA^{Arg}$-TCG | 1 | (Bath *et al.*, 2006; Roine *et al*, 2010) |
| Halfvol4 | 2,109,865-2,163,566 | 2251A-2293A | 53,702 | 14 nt | GGCCTCGGCACTCA | controversial, see main text for details | - | 3 | (Alarcon-Schumacher *et al*, 2022; Hartman *et al*, 2010; Norais *et al*, 2007) |

**Supplementary Table 2. Integration heterogeneity in closely related haloarchaeal genomes at the 14 nt repeat enclosing Halfvol4.**

The genomic regions adjacent to the Halfol4 integration were compared to several closely related haloarchaeal genomes, and integration heterogeneity was observed. For genomes lacking an insert, the position of the empty site (14 nt repeat) is given and insert length is given as "none". In *Haloferax* sp. Atlit-105R the insert is 16 kb and has an unrelated sequence. In *Haloferax alexandrinus strain* pws11 the insert is 61 kb and shows patches of high similarity to Halfvol4 (about 90% sequence identity) which amount to 22 kb.

| | **strain** | **#repeats** | **insert length** | **accession** | **position** |
|---|---|---|---|---|---|
| 1 | *Hfx.volcanii* DS2 | 2 | 53,702 | CP001956.1 | 2109865-2163566 |
| 2 | *Haloferax alexandrinus* strain Wsp1 | 1 | none | CP048738.1 | 2977990-2978003 |
| 3 | *Haloferax lucentense* strain SVX82 | 1 | none | CP104741.1 | 2171542-2171529 |
| 4 | *Haloferax alexandrinus* strain BNX82 | 1 | none | CP106966.1 | 2151664-2151677 |
| 5 | *Haloferax* sp. AS1 | 1 | none | SPSA01000001.1 | 2216069-2216082 |
| 6 | *Haloferax* sp. Atlit-48N | 1 | none | QEQI01000002.1 | 234185-234198 |
| 7 | *Haloferax* sp. Atlit-47N | 1 | none | PSYS01000002.1 | 208273-208260 |
| 8 | *Haloferax* sp. ATCC BAA-644 | 1 | none | AOLF01000021.1 | 282721-282734 |
| 9 | *Haloferax* sp. Atlit-105R | 2 | 16,509 | QPLP01000002.1 | 596510-613018 |
| 10 | *Haloferax alexandrinus* strain pws11 | 2 | 60,967 | WOWC01000001.1 | 2991202-3052168 |

**Supplementary Table 3. Strains, Plasmids and Primers used in this work.**

### A. Strains

| Strain | Genotype | Reference |
|---|---|---|
| **Bacteria** | | |
| E. coli DH5α | F⁻, φ80d*lacZ*ΔM15, Δ(*lacZYA-argF*)U169, *deoR*, *recA*1, *endA*1, *hsdR*17($r_k^-$, $m_k^+$), *phoA*, *supE*44,λ-, *thi*-1, *gyrA*96, *relA*1 | Stratagene |
| **Archaea - *H. volcanii*** | | |
| H26 | DS70 (ΔpHV2), Δ*pyrE2* | (Allers *et al*, 2004) |
| H119 | DS70 (ΔpHV2), Δ*pyrE2*, Δ*leuB*, Δ*trpA* | (Allers *et al.*, 2004) |
| H133 | DS70 (ΔpHV2), Δ*pyrE2*, Δ*leuB*, Δ*trpA*, Δ*hdrB* | (Allers *et al.*, 2004) |
| H100 | DS70 (ΔpHV2), Δ*pyrE2*, Δ*leuB*, Δ*hdrB* | (Allers *et al.*, 2004) |
| ΔHalfvol1-4 | DS70 (ΔpHV2), Δ*pyrE2*, Δ*leuB*, Δ*trpA*, Δ*hdrB*, ΔHalfvol1, ΔHalfvol2, ΔHalfvol3::*trpA*, ΔHalfvol4 | This work |
| ΔHalfvol3 | DS70 (ΔpHV2), Δ*pyrE2*, Δ*leuB*, Δ*trpA*, Δ*hdrB*, ΔHalfvol3::*trpA* | (Alarcon-Schumacher *et al.*, 2022) |
| ΔHalfvol3ΔHalfvol4 | DS70 (ΔpHV2), Δ*pyrE2*, Δ*leuB*, Δ*trpA*, Δ*hdrB*, ΔHalfvol4, ΔHalfvol3:: *trpA* | This work |
| ΔHalfvol1ΔHalfvol3ΔHalfvol4 | Δ*pyrE2*, Δ*leuB*, Δ*trpA*, Δ*hdrB*, ΔHalfvol1, ΔHalfvol4, ΔHalfvol3::*trpA* | This work |
| Δ*oapA* | DS70 (ΔpHV2), Δ*pyrE2, ΔoapA* | (Mills *et al*, 2024; Wolters *et al*, 2019) |

### B. Plasmids

| Plasmid | Characteristic | Reference |
|---|---|---|
| pTA131-UP-DO (Halfvol1) | ColE1 ori, f1 ori, *lacZ*, AmpR , *pyrE2*, Upstream (479 bp) and downstream (500 bp) region of Halfvol1 | This work |
| pTA131-UP-DO (Halfvol2) | ColE1 ori, f1 ori, *lacZ*, AmpR , *pyrE2*, Upstream (761bp) and downstream (740 bp) region of Halfvol2 | This work |
| pTA131-UP-DO (Halfvol4) also termed pTA1102 | ColE1 ori, f1 ori, *lacZ*, AmpR , *pyrE2*, Upstream (3,100 bp) and downstream (1,498 bp) region of Halfvol1 | This work |

### C. Primers

| name | sequence 5´- 3´ |
|---|---|
| US long | GTTGGGTAACGCCAGGGTTTTCC |
| RS long | CAATTTCACACAGGAAACAGCTA |
| M13 | TGTAAAACGACGGCCAG |
| M13r | CAGGAAACAGCTATGAC |
| prov6 circ F | CAGCGGTGAACGTGGTTGTTAG |
| prov6 circ R | CATACACTCAGGGTAGCAAGTG |

| | |
|---|---|
| Prov2 circ R | GTCGAGATACTGCACTACGC |
| Prov2 circ F | GTCAGACGGCTATCCGAGTC |
| prov5 circ F | CTTGACGCACTCGGAGACGAG |
| prov5 circ R | GAGGCGGTGGAACTGAATCAC |
| prov1 circ F | GCAGTACCCGACGAAGTAATC |
| prov1 circ R | GACCTTATCGATTTCAGGAGC |
| Prov6 UP F | CATCGAGCGGCTGGATTGAC |
| Prov6 DO R | GTCCTCGCGGTCATGTTCG |
| UP prov2 F | GTTCCGAACGACGTCAGCATC |
| DO prov2 R | GACTCGCGGGGATATTGAACC |
| DO after tRNA prov2 R | CAAGCGAGTCCAGTGACTGAC |
| Prov5 UP F | GAACTAATCGTCGCGTCCTAC |
| Prov5 DO R | CGAAGTAGTCGTTGATGGCG |
| DO Halfvol4 R | GTTGGAGAAGACGTGAGCGAG |
| DO prov1 R | CTTACTTGTGGAAGAGGCAGAG |
| DO Halfvol4 F2 | GAAAGCCGATGTCGTCGTCG |
| Halfvol4 circ F | GTTCTTCAAGCGTCGACTCCAG |
| Halfvol4 circ R | GCTTTCGGTTGGACGGGACAG |
| circ Halfvol4 F2 | CAATAGGCCTTTACCCGCAC |
| circ Halfvol4 R2 | CTCGGTCCATGTCCAGATAGA |
| HVO_0372 F | GTAGTGTCGGCCGGGACCTC |
| HVO_0372 R | CTGTACAACACCGCACGGCAG |
| pHV4 int F | GAATGGTTATTGACTGGTGC |
| pHV4 int R | CGCAAATCTCGGAGCCTG |
| pHV4 episomal F | GAGCAGTGGTTCTGGCTC |
| pHV4 episomal R | GAATATCTGAGTCTTGCTGAAC |
| ProVir1up.fw | GGACGTCGAGAACGTCTTT |
| ProVir1do.rev | GGCTTTCGACTGCTCGCC |
| ProVir1up.fw | GGACGTCGAGAACGTCTTT |
| ProVir1do.rev | GGCTTTCGACTGCTCGCC |
| HVO2253.int-fw | GGATGGCGCAACAGAATTG |
| HVO2254.int-rev | GGAACAGAGATTCCGTCCTA |
| ProVir1UPSonde-fw | GGGATGGACGAGGTAGGAGT |
| ProVir1UPSonde-rev | TCAACGCCCTGCAGTACAAT |
| Provir1intsonde-fw | AGCTGATTCCGACGTTTCGT |
| Provir1intsonde rev | TCCTCTATCGAAGGGTGCCA |
| ProVir6-up.fw | GTTCTGGTACCGAAAGAAGC |
| ProVir6-up.rev | ATCTCAGTTCGACCGATAG |
| ProVir6-do.fw | CATGCCGACCACCTCAAA |
| ProVir6-do.rev | TCCTCGGCGTGTTCTACT |
| ProVir6-int.fw | GTGGTACCGTTCAGCGAATA |
| ProVir6-int.rev | CGAGTTTGGAGAGTCAGTGG |
| Up KpnI Halvol2 F | TAATATGGTACCCACGAGTCCGGAAGACCTC |

| | |
|---|---|
| Up HindIII Halvol2 R | ATATTAAAGCTTCTCGATTCATCGACAGACAGC |
| Do HindIII Halvol2 F | TAATATAAGCTTCAACACTACCTGCCAGCCG |
| Do XbaI Halvol2 R | ATATTATCTAGACTCTGCGTACGATGAGACTATG |
| HVO_0372 F | GTAGTGTCGGCCGGGACCTC |
| HVO_0372 R | CTGTACAACACCGCACGGCAG |
| CDS1-SnaBI F | TATAATTACGTAATGAGTAGG AATATCTCAGTTCGAC |
| CDS1-XbaI R | AATATTTCTAGATCATGCCGACCACCTCAAACGG |
| CDS2-SnaBI F | TATAATTACGTAATGAGTCACAACACTACCTGCC |
| CDS2-XbaI R | AATATTTCTAGATCATCGCTGGCCCCCGAAATC |
| CDS3-SnaBI F | TAATATTACGTAATGAGTGTTCGACATATCCGAAC |
| CDS3-XbaI R | ATATTATCTAGACTACCATGCACCCCCCTTG |
| qPCR Halfvol2 F1 | TCACAACACTACCTGCCAGC |
| qPCR Halfvol2 R1 | CCGAACGCGAGAAATGGAAC |
| qPCR Halfvol1 F2 | TCATGCCGACCACCTCAAAC |
| qPCR Halfvol1 R2 | AGTGGAGAGTCGGCTACTGAT |
| qPCR Halfvol3 F2 | TATCGACGGCGTAGAGCCAT |
| qPCR Halfvol3 R2 | CATATCCGAACGCGAGAGGT |
